# Phase separation and coexistence in spatial coordination games between microbes

**DOI:** 10.1101/2025.09.09.675176

**Authors:** Guanlin Li, Gabi Steinbach, Peter Yunker, Yao Yao, Joshua S. Weitz

**Affiliations:** Interdisciplinary Graduate Program in Quantitative Biosciences, Georgia Institute of Technology, Atlanta, GA, USA; School of Physics, Georgia Institute of Technology, Atlanta, GA, USA; Department of Biology, University of Maryland, College Park, Maryland, USA; Center for Microbial Dynamics and Infection, Georgia Institute of Technology, Atlanta, GA, USA; Department of Mathematics, National University of Singapore, Singapore; Department of Physics, University of Maryland, College Park, Maryland, USA; University of Maryland Institute for Health Computing, North Bethesda, Maryland, USA

## Abstract

Dense, microbial communities are shaped by local interactions between cells. Both the nature of interactions, spanning antagonistic to cooperative, and the strength of interactions vary between and across microbial species and strains. These local interactions can influence the emergence and maintenance of microbial diversity. However, it remains challenging to link features of local interactions with spatially mediated coexistence dynamics given the significant variation in the microscopic mechanisms involved in cell-to-cell feedback. Here, we explore how microbial interactions over a broad range of ecological contexts spanning antagonism to cooperation can enable coexistence as spatially explicit domains emerge. To do so, we introduce and analyze a family of stochastic coordination games, where individuals do better when playing (*i.e*., interacting) with individuals of the same type than when playing with individuals of a different type. Using this game-theoretic framework, we show that the population dynamics for coordination games is governed by a double-well shaped interaction potential. We find that in a spatial setting this double-well potential induces phase separation, facilitating coexistence. Moreover, we show that for microbes engaged in symmetric coordination games, phase separation takes on a universal scaling that follows ‘Model A’ coarsening, consistent with prior experimental observations for *Vibrio cholerae* mutual killers. Finally, we derive a PDE equivalent of the spatial stochastic game, confirming both the double-well nature of spatial coordination games and the universality of phase separation. Altogether, this work extends prior findings on the link between microbial interactions and population structure and suggests generic mechanisms embedded in local interactions that can enable coexistence.

## I. INTRODUCTION

Bacterial communities are often composed of dense assemblages of cells that include a variety of strains and species. The composition and dynamics of such communities are shaped, in part, by local, cell-to-cell interactions. Microbes have evolved a multitude of mechanisms to interact within and across species – or more generically, among distinct types of microbial cells. Many known mechanisms cause competitive interaction, e.g., some microbes utilize a secretion system that can locally lyse adjacent cells [1, 2]. In contrast, other mechanisms give rise to mutualistic interactions, e.g., by facilitating protection against environmental threats through the production of extracellular barriers or by exchanging resources, including scarce public goods [3–5]. The nature of microbial interactions can also be contingent on context, e.g., turning from competitive to mutualistic as species and strains compete for resources and space [6].

In the context of dense, spatial environments as experienced by aggregates or surface-attached communities, interactions are limited to microbes in close physical proximity. Both the structure and collective dynamics of dense microbial communities arise from locally mediated cell-cell interactions. The community scale structure is an emergent phenomena given the complex feedback dynamics that occur between individual to population scales [7–11]. One common, emergent outcome is local assortment in which distinct microbial types separate into distinct spatial domains comprised of either, but not both, types. This global pattern formation can emerge from initially randomly mixed distributions of cells, stabilize communities, and sustain more diversity than expected in equivalent well-mixed systems [3, 5, 12, 13].

A recent study of strain-mediated killing in *Vibrio cholerae* revealed how spatial ordering and coexistence can emerge in systems dominated by antagonistic interactions [14, 15]. Mechanistically, competing strains mutually kill each other by injecting toxic effectors on contact. Cells express an anti-toxin which renders the toxin ineffectual to cells of the same strain. As a result, initially randomly mixed populations of cells undergo phase separation (‘coarsening’) upon killing and clonal reproduction. Domain formation emerges as cells become locally surrounded by the same type of cell within genetically uniform groups. Numerical simulations of contact-mediated killing demonstrated that phase separation of (nearly) equal competitors exhibits hallmarks of a universal order-disorder transition that is part of the ‘Model A’ (Allen Cahn) class [14] that describes non-conservative systems and was originally developed to describe the interaction of atomic ‘spin’ systems [16].

More generally, the emergent structure and population dynamics of dense microbial communities is a direct result of the local interactions between cells of the same type and between cells of different types. The survival of each strain depends on the ‘strategy’ they utilize, the interaction intensity (the probability of deploying an interaction), and the payoff from the interaction with cells of the same type and of a different type. The problem of how coexistence is linked to individual payoffs is broadly studied in game theory [17] and in the context of spatially explicit systems [18–20]. Evolutionary game theory suggests that symmetries in payoff matrices are sufficient to give rise to similar population dynamic outcomes. Similarly, we hypothesize that mutual killing is one of a subset of microbial interaction mechanisms that can lead to phase separation following the universal Model A transition.

Here, we propose and analyze a game-theoretic framework to model spatial population dynamics that can appear in a broad range of ecological contexts. We do so by investigating a two-player coordination game in which interaction mechanisms can span antagonistic (e.g., mutual killing) to cooperative (e.g., exchange of metabolites or public goods). First, we define transition rules in the (non-spatial) mean-field limit and heuristically derive the replicator dynamics associated with these two-player games – notably we find that these two-player games exhibit alternative stable states, consistent with experimental findings of phase separation. Next, we adapt these transition rules to study phase separation in coordination games in a spatial system. As we show, numerical simulations of spatially explicit two-player coordination games exhibit ‘Model A’ coarsening broadly. These simulations are consistent with experimental findings in the mutual killing case of *V. cholerae* [14, 15] but are not limited to assumptions associated with this particular interaction mechanism. Finally, we derive a reaction-diffusion equation from the hydrodynamic limit of spatial coordination games in continuous space, and show that it resembles the Allen Cahn equation, which describes two-component phase-ordering kinetics in non-conserved systems (Model A class) – suggesting that phase separation of the kind found in antagonistic systems may in fact be a more general feature of dense microbial communities.

## II. A GAME-THEORETIC APPROACH

### 1. Stochastic coordination games

We describe a two-player game where the players represent individual cells of different species that utilize strategies 1 and 2. In this game, a focal player with strategy 1 receives a payoff of *a*_11_ or *a*_12_ when playing against players with strategies 1 (same type) or 2 (different type), respectively. Similarly, focal players with strategy 2 receive a payoff of *a*_21_ or *a*_22_ when playing against players with strategies 1 or 2, respectively (see Fig. 1a). This interaction scenario can be described by a payoff matrix *A* as

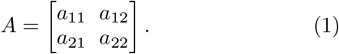

**FIG. 1:**
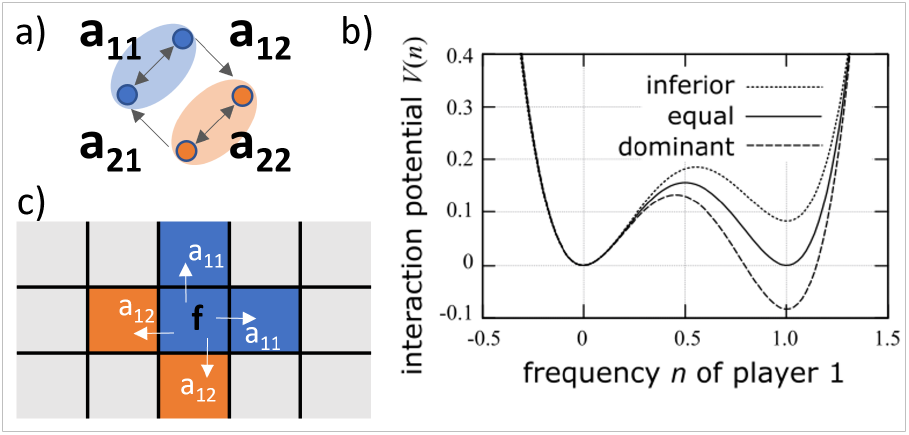
Stochastic Games. a) To characterize a two-species system, we define intra-specific interactions when a focal player with strategy 1 (blue species) or strategy 2 (orange species) plays against their own type, quantified by payoffs *a*_11_ and *a*_22_, respectively, and inter-specific interactions when a player with strategy 1 plays against a player with strategy 2 (payoff *a*_12_), and vice versa (payoff *a*_21_). b) The stochastic game is a deterministic system with an interaction potential *V* (*n*) that is derived from the mean field dynamics of the frequency *n* of player 1 as 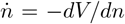 (Eq. 3). For coordination games, *V* (*n*) is a double well potential, as depicted here for three representative coordination games: equally competitive players (competitiveness *C* = (*a*_11_ − *a*_21_)*/*(*a*_22_ − *a*_12_) = 1), and for games where player 1 is dominant (*C >* 1) or inferior (*C <* 1). c) Spatial coordination game on a two-dimensional lattice with focal player *f* with strategy 1 and next neighbor interactions defined by payoff values *a*_*ij*_.

The payoff values, *a*_*ij*_, can be any real positive or negative number. Here, we use the convention that in a biological context, a negative payoff value can lead to death and a positive payoff value can lead to reproduction. The magnitudes of the payoff values are scaled by an effective rate *dt* to obtain birth and death rates. Diagonal elements in the matrix *A* define intra-specific interactions, *i.e*. those that occur between individuals of the same type, and off-diagonal elements define inter-specific interactions that occur between individuals across types. We further focus on the case where the payoff is higher for intra-specific interactions than for inter-specific interactions, *i.e*., *a*_11_ *> a*_21_ and *a*_22_ *> a*_12_. Any game with payoff matrix *A* that obeys these constraints belongs to the class of coordination games.

The community dynamics in the two-player game emerges from the accumulation of birth and death events. Consider a community with *N* members, with a population size *N*_1_ of players with strategy 1 and a population size *N*_2_ of players with strategy 2 such that *N* = *N*_1_ +*N*_2_. We assume a system at carrying capacity where the total community size *N* remains constant at any time. The size *N*_*i*_(*t*) of players with strategy *i* = 1, 2 varies via a continuous-time birth-death process {*N*_*i*_(*t*), *t* ≥ 0}. At any given time *t* ≥ 0, a focal player with strategy *i* plays a game against a randomly chosen opponent with strategy *j*. In the case of positive payoff (*a*_*ij*_ *>* 0), an individual is randomly chosen among the *N −* 1 non-focal members and replaced with an offspring of the focal player at rate *a*_*ij*_; when *a*_*ij*_ *<* 0, the focal player dies and is replaced by a randomly chosen, non-focal individual at rate |*a*_*ij*_|. This convention presumes that predator reproduction can be distinct from consumption of a neighboring prey (as was included to increase the realism of lattice predator-prey models [21]). In the present case, we assume that the lattice occupancy is full, leading to the immediate occupation of an empty site by a randomly selected neighbor, as in [22].

Based on these individual game rules, we now study the community dynamics. In Appendix A, we calculate transition rates between system states for each possible game to heuristically derive the mean field dynamics for the frequency of player 1, *n* := *N*_1_*/N* as *N* → ∞,

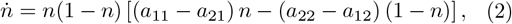

where 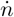 denotes the time derivative of *n*. This is a deterministic system since the strategy of each player is predetermined by the type they belong to upon birth, *i.e*., they cannot change their pre-assigned strategy. Hence, the first-order system in Eq. 2 can be rewritten as a gradient system, 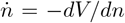, with the potential function *V* (*n*) given as

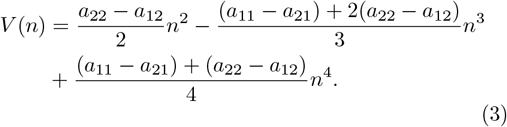

The potential provides insight to the equilibrium configurations of the game; those *n* that fulfill *dV/dn* = 0 correspond to the predictable, steady-state outcomes of the game, where *n*· = 0. We find that coordination games, *i.e*., *a*_11_ *> a*_21_ and *a*_22_ *> a*_12_, lead to a double-well potential (Fig. 1b). It exhibits two local minima of *V* (*n*) at *n* = 0, 1 that correspond to locally stable equilibria, and one local maximum at *n* = *n*_*m*_ ∈ (0, 1) corresponding to an unstable equilibrium, a mixed Nash equilibrium, given by

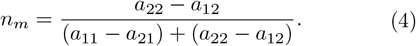

Therefore, the double-well potential *V* (*n*) provides a bistable system, where the outcome of the game depends solely on the initial frequency *n*_0_ of player 1. The population of player 1 goes extinct if *n*_0_ *< n*_*m*_, and it will eliminate the opponent player 2 if *n*_0_ *> n*_*m*_. For equal players, characterized by a competitiveness of *C* = (*a*_11_ − *a*_21_)*/*(*a*_22_ *−a*_12_) = 1, the potential *V* (*n*) is symmetric and *n*_*m*_ = 1*/*2 (Fig. 1b). In contrast, the potential is asymmetric and implies a more shallow well at *n* = 1 with *n*_*m*_ *>* 1*/*2 if player 1 is inferior, i.e., *C <* 1, and a deeper well and *n*_*m*_ *<* 1*/*2 if player 1 is dominant, i.e., *C >* 1.

### 2. Spatial stochastic games

Next, we extend the framework of coordination games to a spatial environment, where the spatial distribution of players and their local environment become relevant. To do so, we apply the stochastic games formalism (see Sect. II 1) on a two-dimensional fully occupied lattice with length *L* per dimension, with a population of *N* = *L*^2^. We apply periodic boundary conditions, *i.e*., the system is a torus in ℤ^2^. Each lattice site *x* ∈ ℤ^2^ is occupied by a player with either strategy 1 or strategy 2. In this spatial model, a focal player can only play with an opponent player that is randomly chosen from its adjacent sites (Fig. 1c); we constrain interactions to the four nearest neighbor sites. Given the strategies of the focal player and the opponent player, the payoff for the focal player is determined by the payoff matrix *A*. For a focal player playing strategy *i* and the opponent playing strategy *j*, the focal player receives payoff *a*_*ij*_ (Fig. 1c). If *a*_*ij*_ *>* 0, then the focal player reproduces at rate *a*_*ij*_ and replaces a randomly chosen adjacent individual with an offspring. If *a*_*ij*_ *<* 0, then the focal player dies with rate |*a*_*ij*_| and is replaced by an offspring of a randomly chosen, adjacent individual. Note that the offspring will always play the same strategy as their parent. At each time step *dt*, players at each lattice site become focal player once to play a game, hence there are *N* games per time step. Complete details of this interacting particle system [23] are given in Appendix D. The pseudo-code of the corresponding Monte Carlo simulation can be found in Appendix B. Next, we discuss the ecological interpretation of the payoff matrix depending on the different combinations of payoff values *a*_*ij*_, before presenting the results of spatial simulations of this interacting particle system in section III.

### 3. Ecological interactions in coordination games

For a two-player game, the sign of *a*_*ij*_ can be interpreted as the ecological impact of player with strategy *j* on player with strategy *i*. Ecological interactions can be defined as either intra-specific (those between individuals of the same type) or inter-specific (those between two types). The use of the terms intra-specific or interspecific interactions usually refer to the ecological interactions between two species instead of two types – however we use the term ‘type’ given our focus on interaction phenotypes here.

Here, we map two-player games to a broad range of intra-/interspecific ecological interactions. The classes of intra-/inter-specific ecological interaction can be characterized by different combinations of the sign of payoff values (Fig. 2). In a two-player system, we define intraspecific interactions in the game as positive if both diagonal elements in matrix *A* have a positive payoff (*a*_*ii*_ *>* 0), mixed if the signs differ (*sgn*(*a*_*ii*_) = −*sgn*(*a*_*jj*_)), and negative if both diagonal elements in matrix *A* are negative (*a*_*ii*_ *<* 0). Further, interspecific interactions are defined as cooperative if off-diagonal elements in matrix *A* are positive (*a*_*ij*_ *>* 0, *i* ≠ *j*), parasitic for mixed signs (*sgn*(*a*_*ij*_) = *−sgn*(*a*_*ji*_), *i* ≠ *j*), and competitive if both off-diagonal elements are negative (*a*_*ij*_ *<* 0, *i* ≠ *j*).

**FIG. 2:**
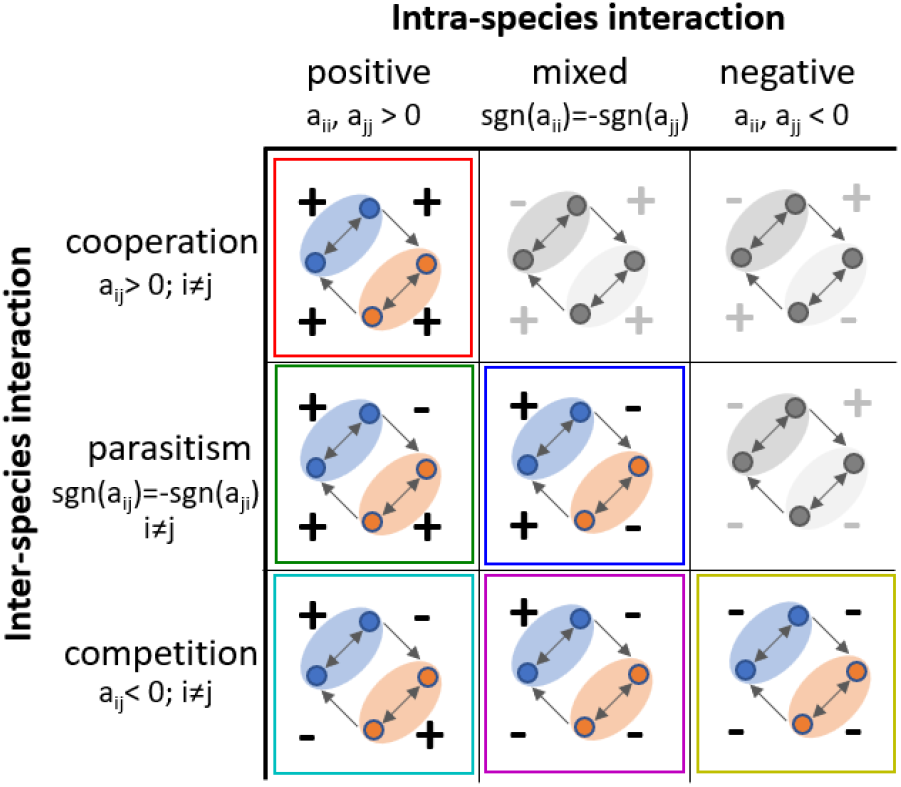
Partitioning games into inter- and intra-specific interactions. Each box corresponds to one class of interaction scenarios, and in analogy to Fig. 1a, the sign (+/−) correspond to payoff values *a*_*ij*_, characterizing positive (+) and negative (−) interactions. For example, the mutual killing system (cyan box) has positive intra-specific interactions, and negative inter-specific interactions. Coordination games exist within the classes that are outlined with a colored box, but not in classes displayed in gray only.

In total, there are four payoff values in the two-player payoff matrix *A*, so there are 2^4^ = 16 possible games (classified by the sign of payoff values, each payoff value *a*_*ij*_ is either positive or negative), see Fig. S1 in Appendix A. In Fig. 2, we summarize all distinct classes of intra-/inter-specific ecological interaction, and highlight those in color that can lead to coordination games.

Based on Fig. 2, we can connect a range of ecological scenarios to a specific class of interactions defined by the sign of payoff values. For example, a competitive scenario of mutual killers (such as for killing-proficient Vibrio strains in [14]) can be modeled via a two-player coordination game with positive diagonal elements and negative off-diagonal elements, which correspond to positive intra-specific interactions and negative (or competitive) inter-specific interactions. Hence, our framework includes a range of mechanistic scenarios including cooperation, parasitism, and competition (*e*.*g*., the mutual killing system).

## III. SPATIAL PATTERN FORMATION

We utilize the spatial coordination game-theoretic framework defined in Sec II 2 to investigate the spatiotemporal dynamics of coordination games in the broad range of ecological contexts shown in Fig. 2. We first model a spatial game with a symmetric payoff matrix with negative off-diagonal elements. We observe rapid phase separation (Fig. 3), similar to the observation for mutual killer biofilms presented in [14]. Domains of players with the same strategy form and grow over time. Next, we determine whether the phase separation undergoes the same class of order-disorder transition as was reported for mutual killers in [14]. We quantitatively characterize spatial organization via the structure factor *S*(*q*) of the spatial dynamics (see Appendix B for more details on spatial processing and calculations). Here, *q* is the wave number coordinate, used to describe spatial structure in wave space after transforming the spatial dynamics from real space to wave space. From *S*(*q*), we obtain the characteristic wave number *q*_*m*_ = ∫ *qS*(*q*)*dq/* ∫ *S*(*q*)*dq*, which represents the effective wave number weighted by *S*(*q*), and serves as a measure for domain size. In Fig. 4, we show the results of spatially explicit simulations with cyan triangles denoting *q*_*m*_(*t*) and *S*(*q*_*m*_) for the coordination game of mutual killers. As a hallmark for ‘Model A’ transitions, *q*_*m*_ scales as *q*_*m*_ ~ *t*^*−*1*/*2^, and *S*(*q*_*m*_) scales as *S*(*q*_*m*_) ~ *t*. Together, this provides a time-independent scaling relationship for 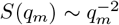, showing that the coordination game of mutual killers undergoes phase separation that follows Model A hallmark scaling.

**FIG. 3:**
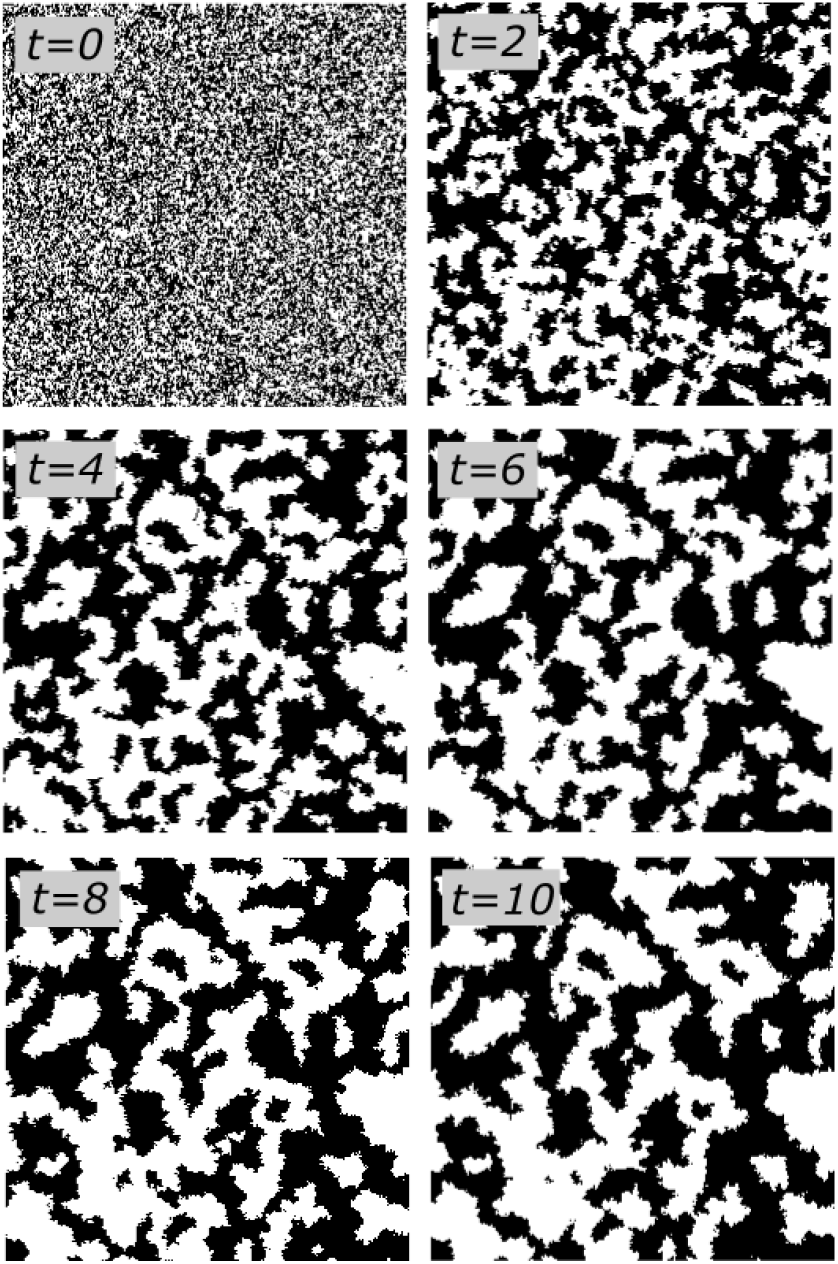
Time lapse visualizations of a two player coordination game with payoff matrix *A* = [10, −5; *−*5, 10]. The images show the spatial configuration (lattice size *L* = 256) at *t* = 0, 2, 4, 6, 8, 10 with dt = 0.05, where the product of payoff values and dt set the magnitude of birth and death probabilities in the Poisson process of game events. White corresponds to player 1 and black is player 2. Starting from well mixed initial condition, initial fraction of player 1 is 0.5, the two players separate into domains whose characteristic length scales (patch size) grow over time.

**FIG. 4:**
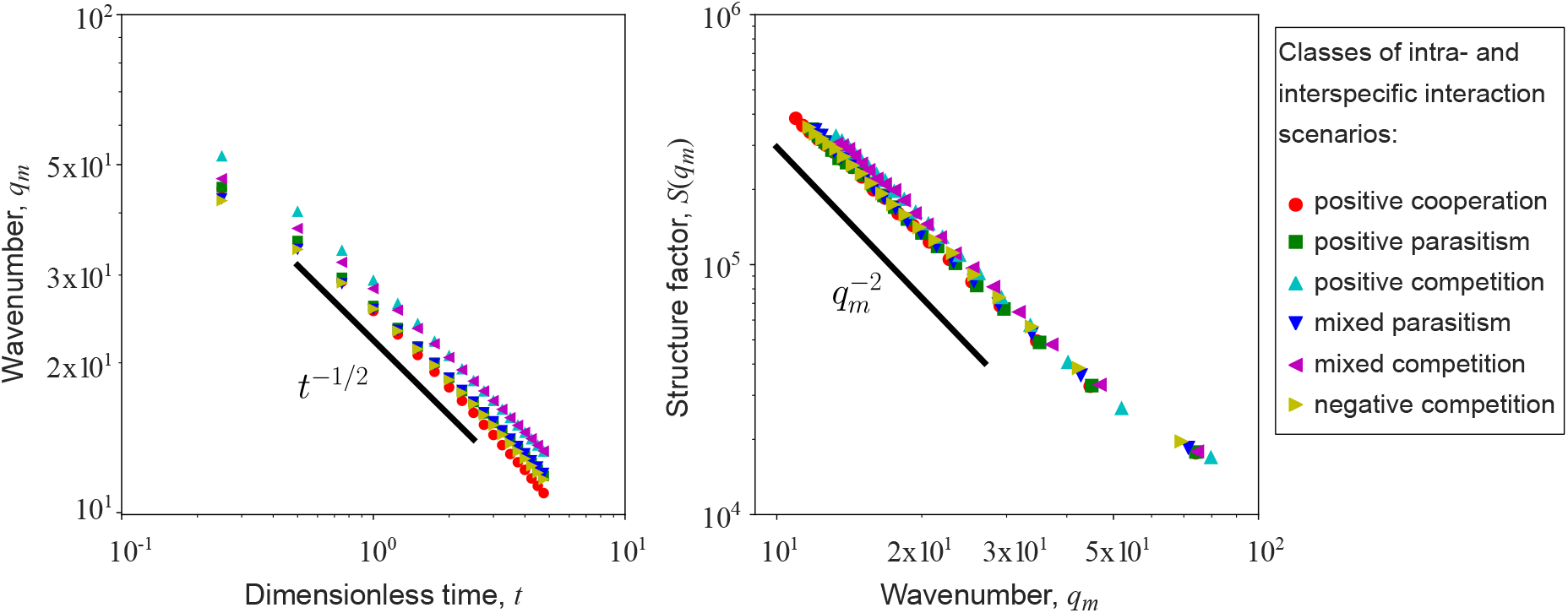
Structural analysis of 6 simulated, symmetric spatial coordination games. In both panels, each curve corresponds to one interaction class in Fig. 2. We chose one example payoff matrix from each class of ecological interactions, every data point is averaged over 50 realizations. The payoff matrices are [10, 2; 5, 7] (red circle), [10, −3; 5, 2] (green square), [2, −1; −3, 4] (cyan triangle), [10, −7; 5, −2] (blue triangle), [2, −7; −3, −2] (magenta triangle) and [−5, −7; −10, −2] (yellow triangle). The relationship between *q*_*m*_ and *t* follow a universal *t*^*−*1*/*2^ trend. Similarly, *S*(*q*_*m*_) curves collapse when *S*(*q*_*m*_) is plotted versus *q*_*m*_ (right), hence, all the games are undergoing the same coarsening transition.

Next, we show that spatial coordination games with equal opponents, characterized by a symmetric double well potential (see Fig. 1b) can lead to ‘Model A’ coarsening. Note that here ‘equal’ means that the competitiveness is *C* = 1, and does not necessarily imply equal payoff values for both players. To do so, we arbitrarily choose a payoff matrix for each class of intra-/inter-specific interaction in Fig. 2 that represents a coordination game, under the constraint of a symmetric double well potential, *i.e*., (*a*_11_ − *a*_21_) = (*a*_22_ − *a*_12_) *>* 0 and a relative difference of 5 between intraspecific and interspecific payoff (Δ*a* = *a*_11_ − *a*_21_ = *a*_22_ − *a*_12_ = 5). We simulate a spatially explicit two-player coordination game for each case and conduct the spatial structural analysis on the resulting spatiotemporal dynamics. We find that for each class of game the system undergoes phase separation. The curves of the structural analysis, *q*_*m*_(*t*) and *S*(*q*_*m*_), collapse for all games (see Fig. 4), and their temporal progression is consistent with the hallmarks of Model A order-disorder phase separation. To demonstrate the universal coarsening dynamics more broadly for symmetric coordination games (*C* = 1), we test the coarsening dynamics for a wider range of values Δ*a* ∈ [1 : 10] of the relative inter/-intraspecific payoff and for a range of intraspecific payoff values *a*_11_, *a*_22_ ∈ [−5 : 5]. We observe that the relationships between *q*_*m*_ and *t*, as well as *S*(*q*_*m*_) versus *q*_*m*_, align with the expected behavior of Model A’s universality more broadly (see Fig. S3 in Appendix C).

In addition, we study the coarsening behavior in spatial coordination games when the double-well potential functions are not perfectly symmetric (Appendix C). Importantly, we observe phase-separation for all investigated classes of coordination games even for asymmetric potentials. We further find that the coarsening behavior of coordination games with a small deviation from symmetric potential (low degree of asymmetry) still closely follows the ‘Model A’ coarsening. In contrast, for coordination games with high degree asymmetric potential, while we can still observe coarsening (domains are formed), the coarsening is no longer consistent with a ‘Model A’ order-disorder phase separation.

## IV. COORDINATION GAMES IN CONTINUOUS SPACE

We extend the prior analysis of interacting particle systems to investigate coarsening dynamics within two-player games in continuous space. In continuous space, phase separation processes that follow ‘Model A’ order-disorder transition can be described by a reaction–diffusion equation, the Allen-Cahn equation [16]. In the present context, the reaction term emerges from the same interaction between opponents as defined by the game rules applied on a lattice in the previous section. However, here we consider a scaled lattice 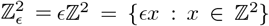, where is the lattice constant. Further, the frequency of types in the particle system are represented as expected means, with *u*^*E*^(*t, x*) being the frequency of player 1 at time *t* and position *x*, and *v*^*E*^(*t, x*) = 1 − *u*^*E*^(*t, x*) the frequency of player 2. Based on the spatial game rules, we identify flip rates for the state of lattice sites in Appendix D, and use them to derive the reaction term *f* (*u*) in Appendix E. Next, we need to extend our model to include a diffusion term, which emerges from cellular migration. Since the lattice is fully occupied, diffusion is realized when individual players swap positions with one of their nearest players. We define that players at neighboring sites will be allowed to exchange sites (perform random walks) at rate 4^*−*2^ on a lattice with spacing.

With these ingredients, we can establish the partial differential equation (PDE) limits of spatial coordination games. In the hydrodynamic limit of the interacting particle system, in which the lattice spacing goes to zero (ϵ→0) and the speed of stirring goes to infinity (ϵ^*−*2^→ ∞), the frequencies of the two types of particles converge to continuous solutions of a reaction-diffusion equation. We can write the time derivative of the population *u* of player 1 as:

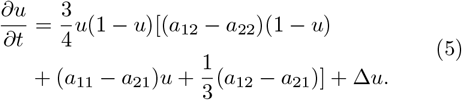

The detailed derivation of Eq. 5 can be found in Appendix E. The central limit theorem suggests that in the limit players perform Brownian motion (by appropriate scaling). Notably, the reaction term in Eq. 5 differs from the mean-field replicator dynamics in Eq. 2. Hence, the limiting reaction-diffusion equation associated with the spatial game system is different from simply adding a diffusion term to the mean-field replicator dynamics. This observation recapitulates a central observation of earlier studies that explicitly derived reaction-diffusion models for interacting particle systems [24]. The difference occurs as for each spatial game in dimension 2, there is a 1/4 probability that the random third player selected as the site of a potential birth event coincides with the site of the opponent – this probability is not negligible. In contrast, in a non-spatial (mean-field) system with *N* players the probability would be 1*/N*, which goes to zero as *N* → ∞. Notably, the replicator dynamics can be recovered in a PDE system with large dimension *d* or by modifying the game rules such that the opponent player cannot be identified as the randomly selected third player (see Appendix E, Remark 1, for more details).

Finally, we examine the solution space of the PDE model. The reaction term in Eq. 5 is a cubic polynomial. In analogy to section II 1, we define a potential function *W* as

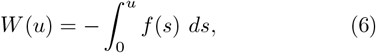

where *f* is the reaction term in Eq. 5. If *W* has two local minima at *u* = 0 and *u* = 1, and a local maximum at *u* = *u*_*m*_ where *u*_*m*_ ∈ (0, 1), then *W* is a double-well potential (see Appendix E and E9 specifically for the condition of *W* being a double-well potential). As a result the reaction term will induce phase separation.

To consider the impact of the diffusion term on coarsening, we need to examine the complete reaction-diffusion equation in Eq. 5, which takes the form of an Allen-Cahn equation [16]. When Eq. 5 is defined on a two dimensional torus 𝕋^2^ (periodic boundary condition), the evolution of Eq. 5 can be viewed as the *L*^2^-gradient flow of the Ginzburg-Landau free energy functional [25, 26]

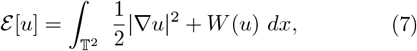

*i.e*., 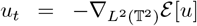. The free energy is nonincreasing in time (see Appendix E). The free energy ℰ [*u*] contains two terms; the gradient term penalizes spatial variation (thus, diffusion has a smoothing effect), while the potential *W* (which has a double-well shape with two local minima) drives the system to undergo phase separation. This conclusion is supported by the linear stability analysis of a mixed-strategy homogeneous steady state *ū*, see Appendix E.

## V. DISCUSSION

Microbes exhibit a multitude of interaction mechanisms that shape the structure and dynamics of microbial communities. In this work, we have used a game theoretic framework to systematically evaluate microbial spatiotemporal dynamics given a range of ecologically relevant combinations of inter- and intra-specific interactions. We find that in the case of coordination games, where cells of the same microbial type prefer to coordinate with kin over non-kin, the game rules always lead to a bistable interaction potential. In well-mixed environments this leads one type to win and exclude the other type. In contrast, when applying this game in an individual-based spatial setting, where interactions are restricted to nearest neighbors, we observe coarsening of the ‘Model A’ universality class in all studied classes of coordination games, facilitating coexistence between types. Our results demonstrate that comparable population dynamics can emerge irrespective of the underlying type of interaction, providing a path forward to connect local interaction rules with the structure and dynamics of complex microbial communities[1, 3, 6, 9, 11, 14, 27, 28]. As we show, phase separation occurs in spatially interacting coordination games for a range of interaction types and strengths. Previously, phase separation was shown to emerge in the specific case of two-player antagonistic scenarios, and universal behavior is obtained for symmetric antagonism [14]. We find that the observed universality class is more general: ‘Model A’ coarsening is obtained independent of the sign of ecological interaction spanning antagonistic to cooperative insofar as the payoff matrix corresponds to a symmetric potential. Moreover, by extending the spatial analysis of the coordination game to a continuous-space model, we demonstrated that the coarsening dynamics driven by the double-well interaction potential is captured by a non-conserving reaction-diffusion equation that represents the hydrodynamic limit of spatial games. In doing so, we find that the reaction-diffusion equation associated with the spatial game system is different from one obtained by adding a diffusion term to the replicator dynamics (i.e. to the mean-field limit of the corresponding non-spatial system). This observation recapitulates the findings in [24] and reinforces the importance of assessing how microscopic spatial interactions transform the emergence of spatiotemporal patterns in biological systems.

These findings come with caveats. The structure of our mean-field and spatially explicit analysis assumes dense communities with constant total population sizes. In reality, both the frequency of types and the total number of cells are dynamic, e.g., because of exogeneous changes in resource availability and due to endogenous feedback. Extending the current model to include coupling between game and environment could lead to novel dynamics [22, 29–31]. Similarly, the present framework focuses on types with fixed interaction traits. As a result, the emergent spatiotemporal dynamics are limited to the study of ecological interactions rather than the broader problem set of eco-evolutionary dynamics [32]. Intriguingly, the emergence of domains of single types of microbes could shape selection and subsequent evolution. As the frequency of contacts between kin increases and the contacts between non-kin decreases upon phase separation, the switch in social contexts may give way to selection pressure both for functional traits that contribute to initial phase separation and the formation of clonal domains and for functional traits which stabilize locally similar, microbial domains [33]. Finally, while our work focuses on pairwise interactions only, it it can be readily expanded to study many-player systems to explore generic mechanisms that may underlie emergent coexistence in multi-species systems [34] and the role of interaction strength [8, 35] to promote diversity.

In sum, our work demonstrates generic mechanisms that can drive two-player systems to universal modes of coexistence over a range of ecological scenarios, inspired by - but not limited to - experimental findings of phase separation within mutual killing strains of *V. cholerae*. In doing so, we have shown how to formally connect features of local ecological interactions with emergent patterns at population scales. Although limited in its focus on static traits without evolution or environmental feedback, we hope that the explicit treatment of stochastic spatial games can provide a basis for useful extensions of interest to a broader range of microbial systems and contexts.

## ACKNOWLEDGMENTS

JSW acknowledges support from the Army Research Office (W911NF1910384), the Simons Foundation (930382), and the Chaire Pascal program of the Ile-de-France region. JSW is an investigator at the University of Maryland-Institute for Health Computing, which is supported by funding from Montgomery County, Maryland and The University of Maryland Strategic Partnership: MPowering the State, a formal collaboration between the University of Maryland, College Park and the University of Maryland, Baltimore. P.J.Y. acknowledges funding from the NIH National Institute of General Medical Sciences (grant no. 1R35GM138354-01). YY acknowledges support from the NUS start-up grant, MOE Tier-1 grant, and the Asian Young Scientist Fellowship.

## DATA AVAILABILITY

All code for simulations and plotting is available at https://github.com/Gabi-Steinbach/Spatial-microbial-games and archived at https://doi.org/10.5281/zenodo.17039081

## Appendix A

**Derivation of mean-field limit**

Throughout the appendix, we denote a player with strategy 1 by ➀, and a player with strategy 2 by ➁. Using the notation of chemical reactions [36], we list below all the possible games that result in a change in the state *N*_1_. If the randomly selected third player is not the opponent, the games that result in a change in the state *N*_1_ are as follows:

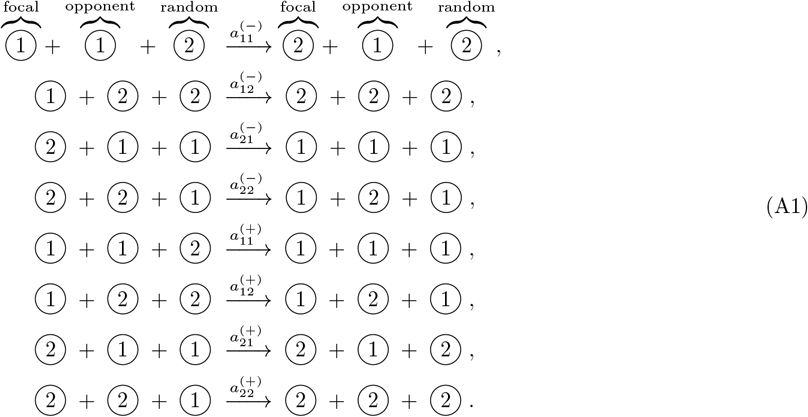

Here the reaction rates 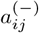 and 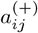 are shorthands of 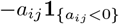 and 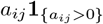, where **1**_*{E}*_ is the indicator function; **1**_*{E}*_ = 1 if event *E* is true and **1**_*{E}*_ = 0 otherwise. Technically, for any given payoff matrix *A* ∈ ℝ2*×*2, only four reactions in Eq. A1 are realized. If the randomly selected third player coincides with the opponent, the following games would result in a change in the state *N*_1_:

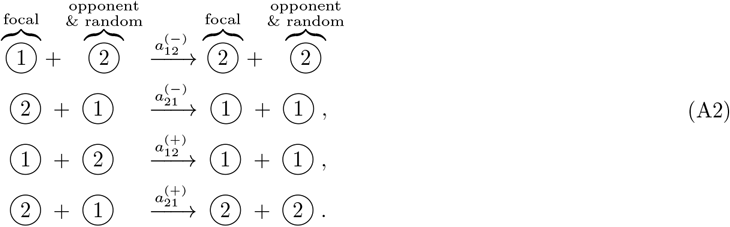

Next, we derive the mean-field limit of above stochastic games by following a similar exercise in [22]. Taking into account all the cases in Eq. A1 and Eq. A2, we can write the transition rates as

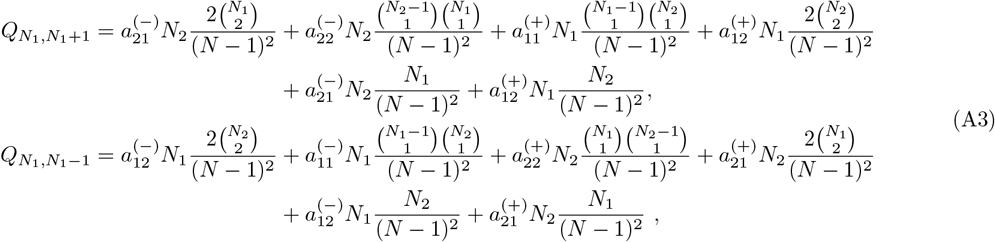

where *Q*_*n,n*′_ represents the transition rate from state *n* to state *n*′. Note that *Q*_*n,n*′_ = 0 if |*n − n*′ | *>* 1. For each equation in Eq. A3, the first four terms on the right hand sides correspond to the cases in Eq. A1, whereas the last two terms correspond to the cases in Eq. A2. The evolution of state *N*_1_ is governed by the master equation, which has a gain-loss form [22, 39]

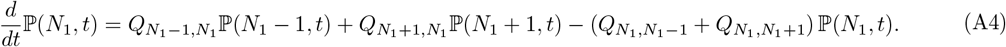

Then, multiplying Eq. A4 with *N*_1_ and summing over *N*_1_, we obtain

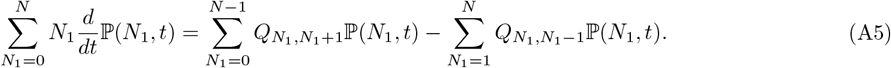

Thus, for the average of the stochastic variable *N*_1_(*t*), its time evolution satisfies

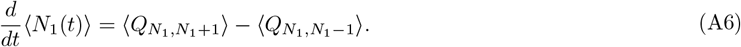

Assume *N, N*_1_, *N*_2_ ≫ 1, we note that

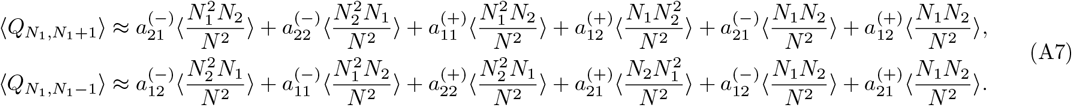

The approximation error can be estimated by calculating the relative change between the components in Eq. A7 and corresponding components in Eq. A3. For example, we compute the relative error for the first component as follows:

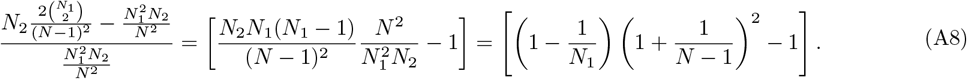

This error is on the order max(1*/N*, 1*/N*_1_). As *N, N*_1_, *N*_2_ 1, the relative error approaches zero. Here, we define *n* = *N*_1_*/N*, and thus *N*_2_*/N* = 1 − *n*. Plugging the expressions of averaged transition rates in Eq. A7 into Eq. A6, then we normalize the system via dividing both sides of Eq. A6 by *N*, we find

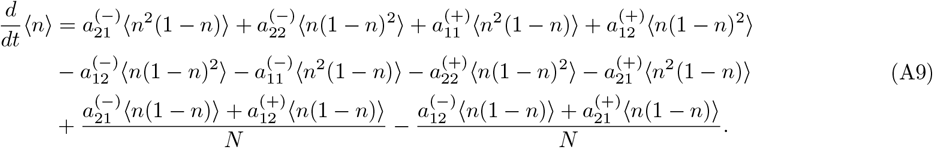

Since *N* ≫ 1, the last two terms in the order of 1*/N* are dropped, and Eq. A9 becomes

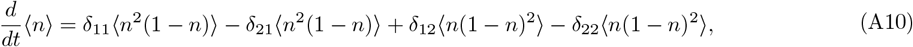

where 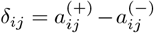. Note that 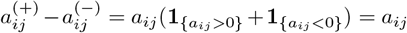. In the limit of large *N*, the normalized system converges to the deterministic dynamics, *i.e*., the fluctuations around the average value *n* are negligible. We thus omit the angular brackets in Eq. A10, and obtain

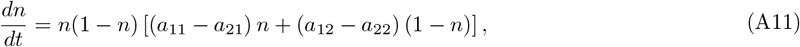

which recovers the standard replicator dynamics.

The details of mapping from two-player games to intra-/inter-specific ecological interactions are shown in Fig. S1.

**FIG. S1:**
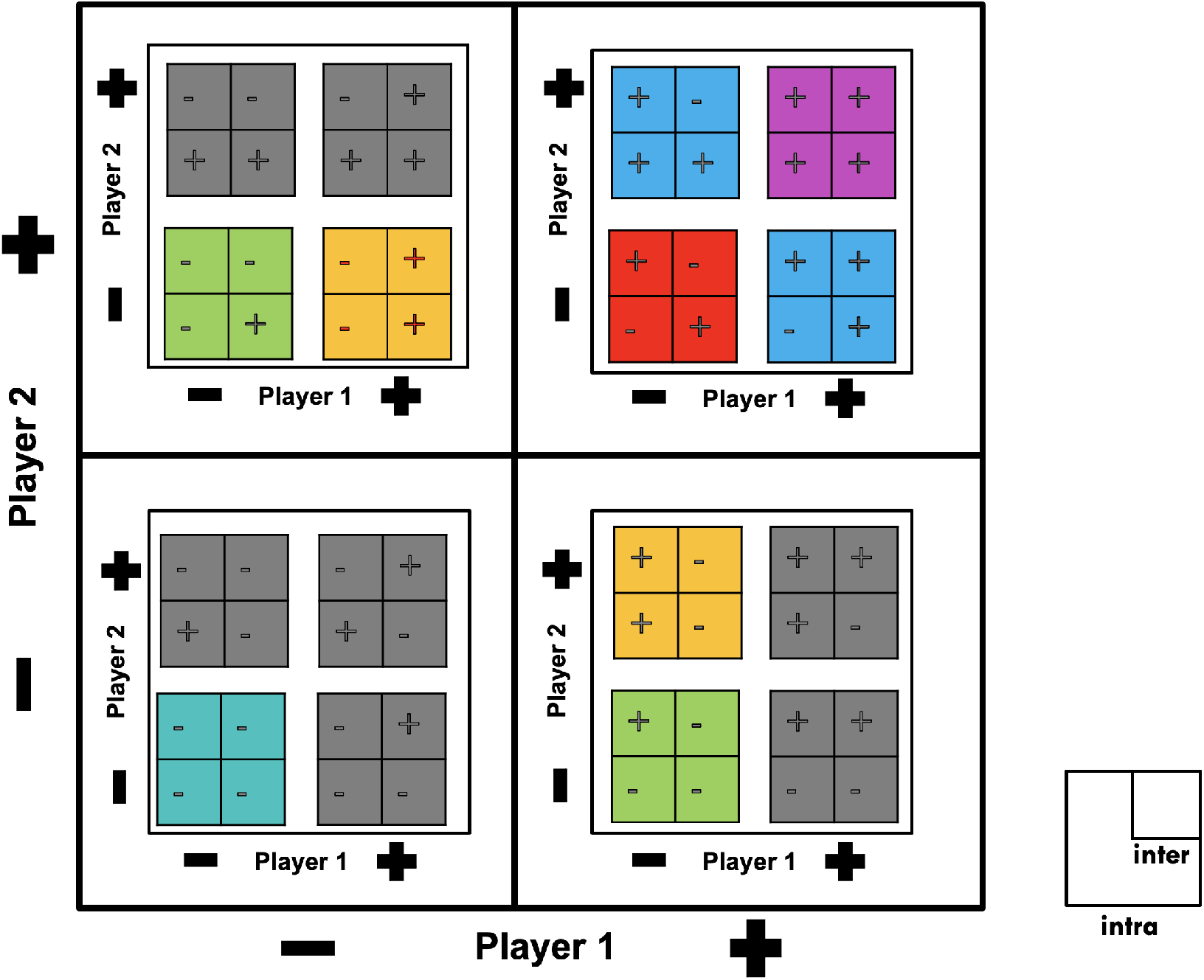
Mapping from two-player games to inter- and intra-specific interactions. The sign (+/−) of payoff values correspond to the positive (+) and negative (−) interactions. There are two layers, the outer-layer represents the intraspecific interactions and the inner-layer represents the inter-specific interactions. Some intra-/inter-specific ecological interactions cannot be mapped to any coordination game (gray blocks), e.g., an example of *anti-coordination game* with sign {*A*} = [−, +; −, +]. Some pairs of games are symmetric (blocks with same color), *i.e*., one can be transformed to another via swapping player 1 and player 2. Hence, there are six distinct inter-/intra-specific ecological interactions that are mapped to coordination games as shown in Fig. 2 in main text.

## Appendix B

**Simulation of spatial games and spatial structure analysis**

The simulation details of spatial games are summarized in Algorithm 1.

### Algorithm 1

Simulation protocol of spatial games

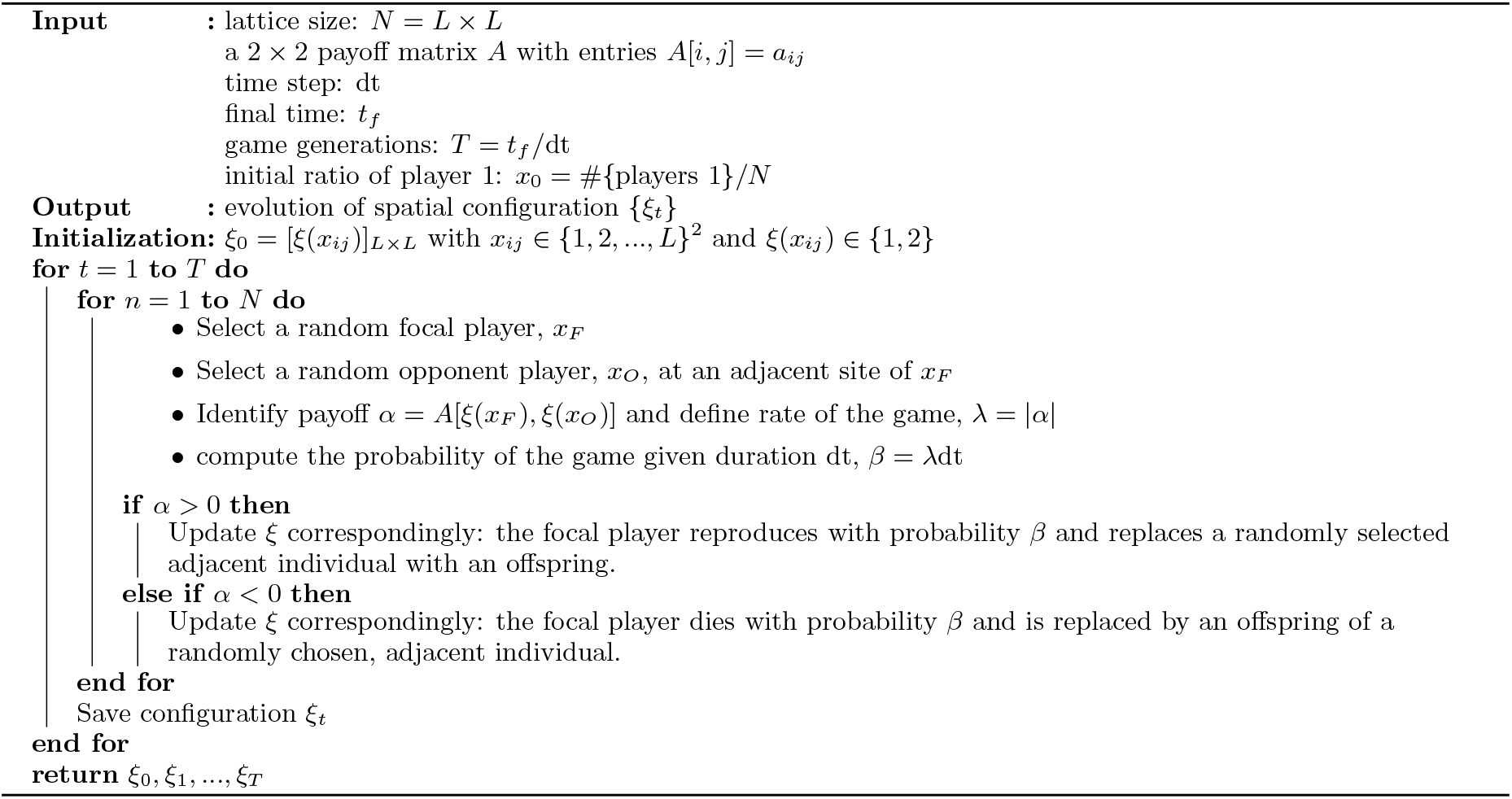

We quantitatively examine the dynamics of phase separation via spatial structural analysis as follows. We first compute the Fourier transform of the spatial configurations obtained from simulation (Figs. S2a, b). This transforms the image from real space into wave space with wave number coordinates *q* = [*q*_*x*_, *q*_*y*_]^*T*^, providing the spatial frequencies in the *x* and *y* directions, respectively. Next, we calculate the structure factor *S*(*q*) by radially integrating the real part of the Fourier transformed image (Fig. S2c) and calculate the characteristic wavenumber as *q*_*m*_ = ∫*qS*(*q*)*dq/* ∫*S*(*q*)*dq*.

**FIG. S2:**
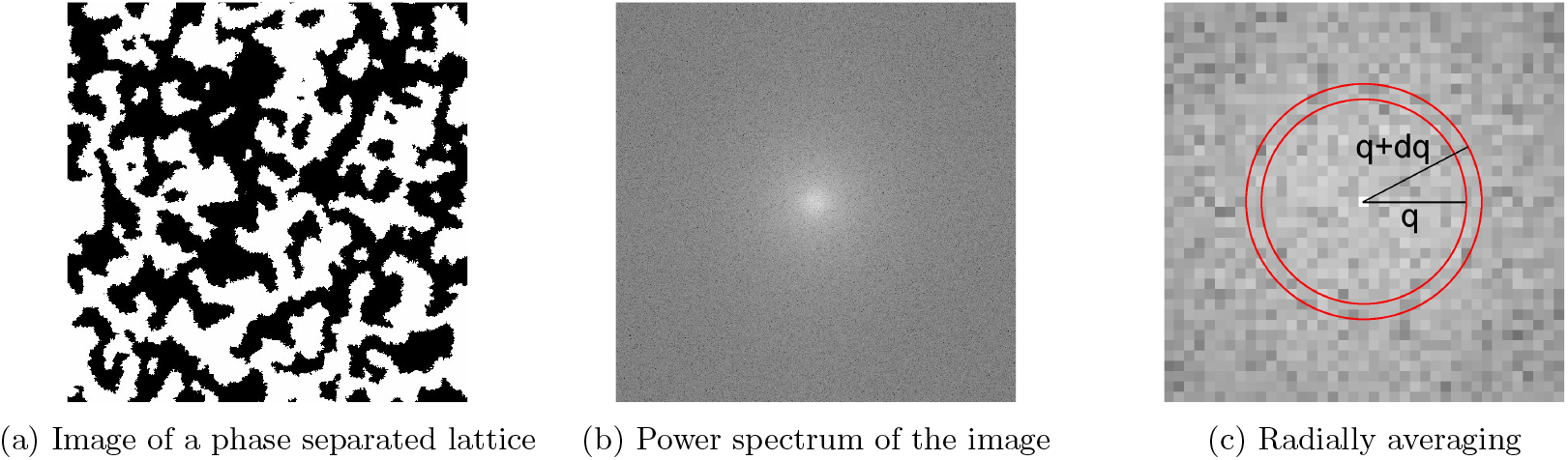
Workflow of the image processing and analysis. a) Black-white image of a phase separated lattice, and b) the power spectrum of the Fourier transformed image in (a), displayed with logarithmic scaling. c) Magnified cutout of the center of the Fourier transformed image (b) illustrating the radial integration for calculating the structure factor *S*(*q*). Intensities of pixels between concentric circles of *q* and *q* + *dq* (red) are integrated and normalized by the number of pixels in this ring. This procedure is performed for the full range of *q*-values to provide the structure factor *S*(*q*) as a function of distance *q* from the center of the Fourier transformed image. Note that the radial axis is displayed with a logarithmic scale, and the image center corresponds to *q* = 0.

## Appendix C

**Coarsening in spatial coordination games: numerical simulations**

In the main text, ‘Model A’ coarsening has been observed for each of the six spatial coordination games with a symmetric double-well potential (competitiveness ratio *C* = 1) and a relative difference between intraspecific and interspecific payoff of 5 (Fig. 4). To more broadly analyze the universal coarsening dynamics for symmetric coordination games, we test a wider range of parameter values. We observe that the relationships between *q*_*m*_ and *t*, as well as *S*(*q*_*m*_) versus *q*_*m*_, align with the expected behavior of Model A’s universality more broadly, as shown in Fig. S3.

**FIG. S3:**
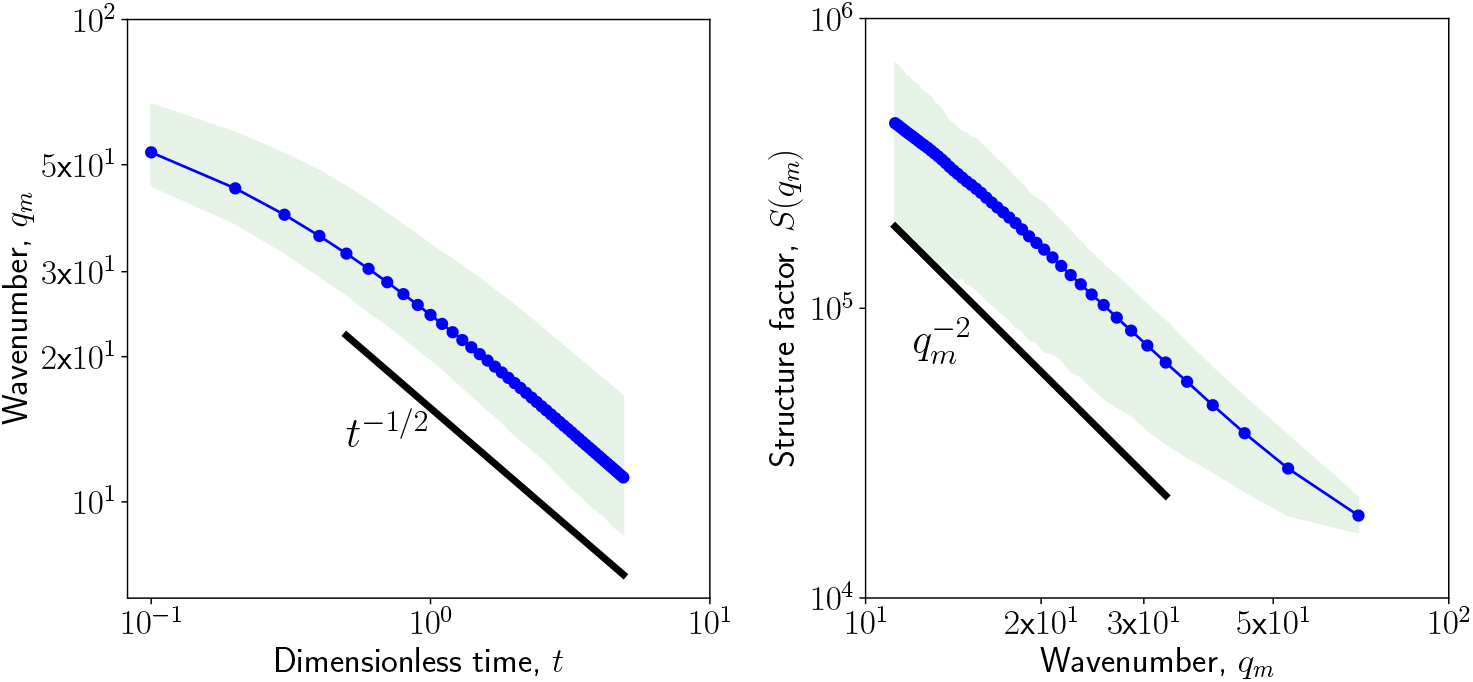
Structural analysis of spatial coordination games with symmetric double-well potential across a broader range of parameter values. Integer values for *a*_11_ and *a*_22_ are independently selected within the range of −5 to 5. The relative difference between intraspecific and interspecific payoff (Δ*a* = *a*_11_ − *a*_21_ = *a*_22_ − *a*_12_) is chosen to range between 1 and 10, leading to the relationships *a*_21_ = *a*_11_ + Δ*a* and *a*_21_ = *a*_22_ + Δ*a*. Simulations are performed for 1,000 payoff matrices (=10×10×10, excluding zero values). For each matrix, we examine the structural relationships between *q*_*m*_ and dimensionless time *t* and between *S*(*q*_*m*_) and *q*_*m*_, averaging over 50 realizations. The averaged curves (blue line) are plotted within the confidence interval band (green shaded area). The 95% confidence intervals for these relationships, across the tested parameters, are consistent with the expected behavior of Model A’s universality.

In analogy to the study of symmetric coordination games, next, we consider two types of asymmetric coordination games: those with a small and a large deviation from a symmetric potential. In the main text Fig. 4, we choose one example payoff matrix with symmetric double-well potential from each class of ecological interaction. The payoff matrices are [10, 2; 5, 7], [10, −3; 5, 2], [10, −7; 5, −2], [2, −1; −3, 4], [2, −7; −3, −2] and [−5, −7; −10, −2]. The relative fitness values (or the intra-/inter-specific interaction payoff differences) of the two players are the same for all studied examples, *i.e*., (*a*_11_ − *a*_21_) = (*a*_22_ − *a*_12_) = 5. For convenience, we consider these payoff matrices as our baseline cases. To obtain asymmetric potentials, we add a deviation factor *η* to control the degree of asymmetry. For example, adding *η* to the payoff matrix *A* = [10, 2; 5, 7] with symmetric potential yields *A*_*η*_ = [10, 2; 5 −*η*, 7]. In our numerical experiments and coarsening analysis, we use *η* = 0.5 for a low degree of asymmetry and *η* = 3 for a high degree of asymmetry.

The coarsening behavior of coordination games with a low asymmetry potential still closely follows the ‘Model A’ coarsening, see Fig. S4. For coordination games with high degree of asymmetrical potential, we can still observe coarsening behavior, see Fig. S5a, however, the coarsening is not consistent with ‘Model A’ order-disorder phase separation, see Fig. S5b.

**FIG. S4:**
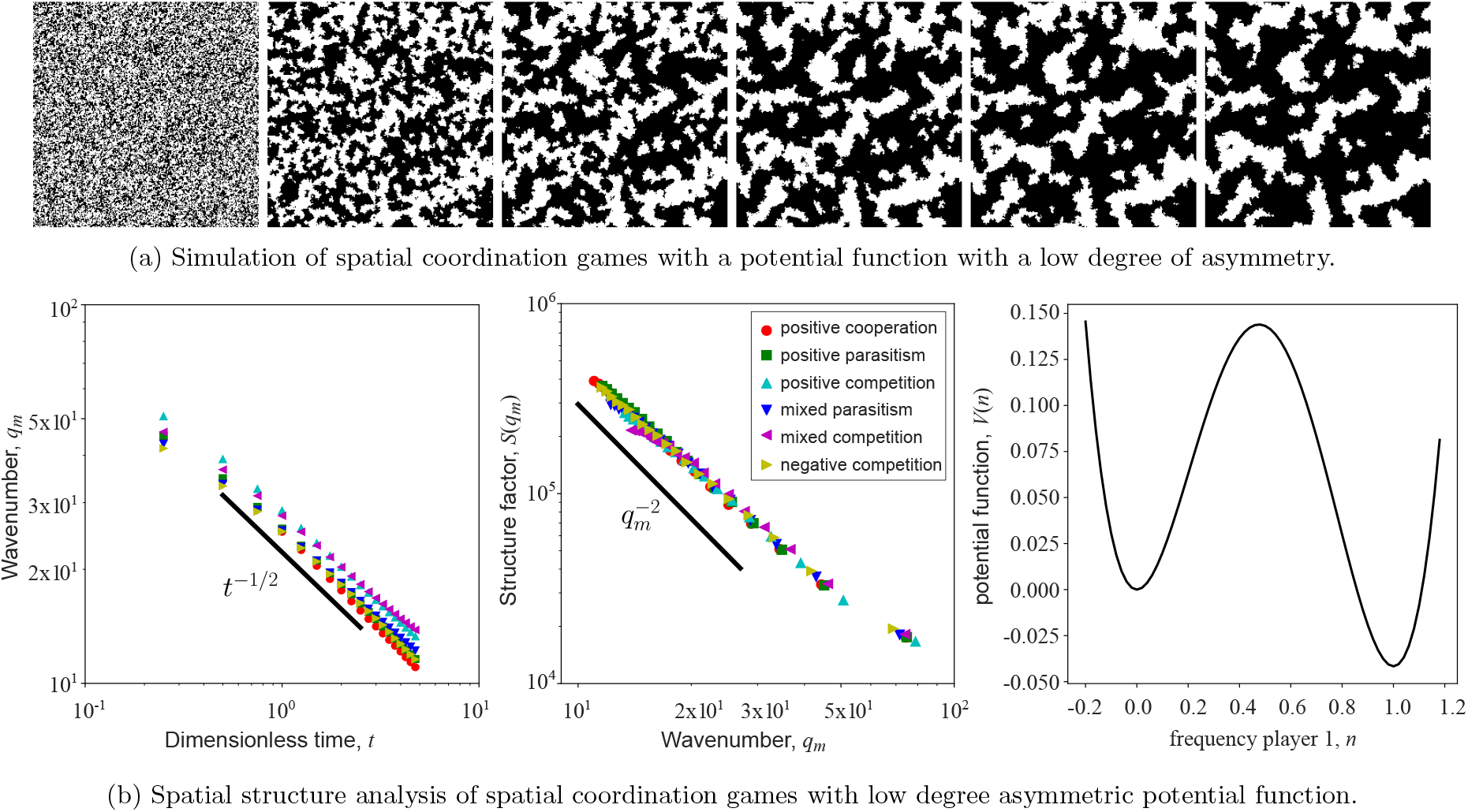
Spatial coordination games with a low degree of asymmetry in the payoff matrix. (a) Time lapse images of a two player coordination game with payoff matrix *A* = [10, −5; −5.5, 10]. The images show the spatial configuration (lattice size *L* = 256) at *t* = 0, 2, 4, 6, 8, 10 with dt = 0.05. White corresponds to player 1 and black is player 2. Starting from a well mixed initial condition with an initial fraction of 0.49 for player 1 (unstable equilibrium of corresponding replicator equation), the two players separate into domains which grow over time. (b) Structural analysis of simulated spatial coordination games that correspond to asymmetric potentials with a small deviation (*η* = 0.5). For both panels (left and middle), there are 6 different colors, each corresponding to one type of interaction class in Fig. 2. We choose one example payoff matrix from each class, and every data point is averaged over 50 realizations. The payoff matrices are [10, 2; 4.5, 7] (red circle), [10, −3; 4.5, 2] (green square), [10, −7; 4.5, −2] (blue triangle), [2, −1; −3.5, 4] (cyan triangle), [2, −7; −3.5, *−*2] (magenta triangle) and [−5, −7; −10.5, −2] (yellow triangle). The relationship between *q*_*m*_ and *t* is summarized in the left panel; all games closely follow a universal *t*^*−*1*/*2^ trend. Similarly, *S*(*q*_*m*_) curves collapse when *S*(*q*_*m*_) is plotted versus *q*_*m*_, all the games undergo the same coarsening process, see middle panel. The plot of the potential function on the right panel shows the low degree of asymmetry.

**FIG. S5:**
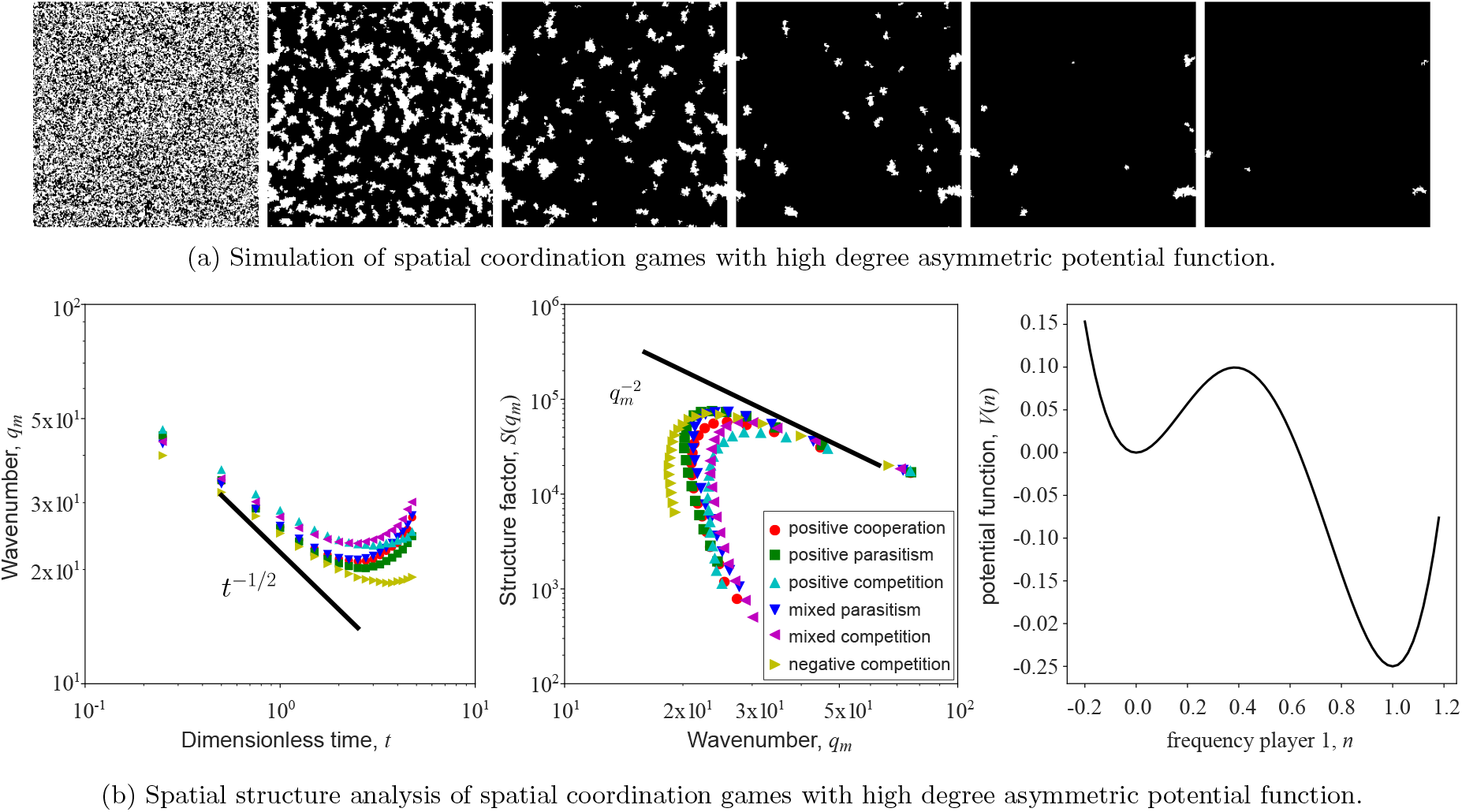
Spatial coordination games with a high degree of asymmetry in the payoff matrix. (a) Time lapse images of a two player coordination game with payoff matrix *A* = [10, −5; −8, 10]. The images show the spatial configuration (lattice size *L* = 256) at *t* = 0, 2, 4, 6, 8, 10 with dt = 0.05. White corresponds to player 1 and black is player 2. Starting from a well mixed initial condition with an initial fraction of 0.45 for player 1 (unstable equilibrium of corresponding replicator equation), the two players separate into domains which grow over time. (b) Structural analysis of simulated spatial coordination games that correspond to asymmetric potentials with a large deviation (*η* = 3). For both panels (left and middle), there are 6 different colors, each corresponding to one type of interaction class in Fig. 2. We choose one example payoff matrix from each class of interaction, and every data point is averaged over 50 realizations. The payoff matrices are [10, 2; 2, 7] (red circle), [10, −3; 2, 2] (green square), [10, −7; 2, −2] (blue triangle), [2, −1; −6, 4] (cyan triangle), [2, −7; −6, −2] (magenta triangle) and [−5, −7; −13, −2] (yellow triangle). The relationship between *q*_*m*_ and *t* is summarized in the left panel; the games deviate from a universal *t*^*−*1*/*2^ trend. Similarly, *S*(*q*_*m*_) curves do not collapse when *S*(*q*_*m*_) is plotted versus *q*_*m*_, see middle panel. The plot of the potential function on the right panel shows the high degree of asymmetry.

## Appendix D

**Theoretical framework of spatial games, interacting particle systems**

The spatial interactions of players can be modeled using the framework of interacting particle systems [23]. Here, we consider a lattice with *L* sites per dimension with periodic boundary conditions. In two dimensions, this is a torus in ℤ^2^ with *N* = *L*^2^ lattice sites, where each site *x* is occupied by a player with either strategy 1 or strategy 2. Hence, the state space of each site is 𝒜 = {1, 2}, and the configuration of the system at time *t* is *ξ*_*t*_ : ℤ^2^ → 𝒜. Let us define the interaction neighborhood of site *x* as

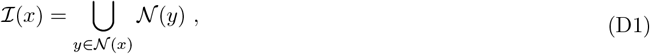

where 𝒩 (*x*) is the set of 4 nearest neighbors of *x*: 𝒩 (*x*) = {*y* ∈ ℤ^2^ : ‖*y x*‖ _1_ = 1}. Only one event is allowed at a time, and the other sites remain (temporarily) unchanged, *i.e*., a regular Markov jump process. We define *c*_*ij*_(*x, ξ*) as the rate at which site *x* at state *i* ∈ 𝒜 flips to state *j* ∈ 𝒜. Note that the flip rate *c*_*ij*_(*x, ξ*) only depends on the configuration *ξ* in the interaction neighborhood ℐ (*x*) of site *x*. We refer to this flip event as the particle flip dynamics. The focal player’s position *x* is selected at random, and the opponent’s position is chosen randomly from the nearest neighborhood (*x*). As we apply the game rules in Eq. A1, the third player’s position is randomly chosen from (*x*) as well. Note that the opponent in a game could be chosen as the third player with some nonnegligible probability 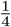 (in contrast to the mean field limit of non-spatial games). For convenience, we define the translation of a site *x* as

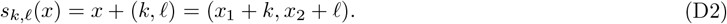

We denote the positions of the focal player, the opponent, and the randomly chosen third player as *x*_*F*_, *x*_*O*_ and *x*_*R*_ respectively. We describe the spatial dynamics by listing all the possible ‘local’ games that may result in a state flip of site *x*.

First, we assume *ξ*(*x*) = 1, the position of the focal player being *x*, and discuss the following scenarios.

- The position of the focal player is *x, i*.*e*., *x*_*F*_ = *x*, the opponent is from 𝒩 (*x*) and it has the same strategy as the focal player, *i.e*., *ξ*(*x*_*O*_) = *ξ*(*x*_*F*_) = 1. Suppose the focal player loses a game and gets replaced by a randomly chosen third player in 𝒩 (*x*) (other than the opponent) with a different strategy, *i.e*., *ξ*(*x*_*R*_) = 2. So, we require *ξ*(*x*) = 1, *ξ*(*x*_*O*_) = 1 and *ξ*(*x*_*R*_) = 2. With *x*_*F*_ = *x* fixed, there are 12 feasible combinations of positions *{x*_*F*_, *x*_*O*_, *x*_*R*_*}*. For every feasible combination, say *{x, s*_1,0_(*x*), *s*_0,1_(*x*)*}*, the reaction (flip dynamics) is

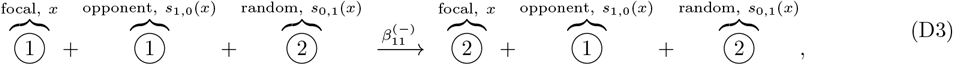

where the flip rate 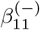 is

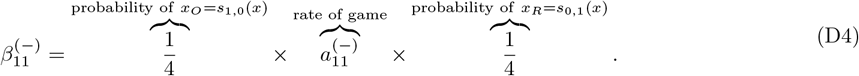
- The position of the focal player is *x, i*.*e*., *x*_*F*_ = *x*, the opponent is from 𝒩 (*x*) and it has a different strategy from the focal player, *i.e*., *ξ*(*x*_*O*_) = 2. Suppose the focal player loses a game and gets replaced by a randomly chosen third player (other than the opponent) with same strategy as the opponent, *i.e*., *ξ*(*x*_*R*_) = 2. So, we require *ξ*(*x*) = 1, *ξ*(*x*_*O*_) = 2 and *ξ*(*x*_*R*_) = 2. With *x*_*F*_ = *x* fixed, there are 12 feasible combinations of positions *{x*_*F*_, *x*_*O*_, *x*_*R*_*}*. For every feasible combination, say *{x, s*_1,0_(*x*), *s*_0,1_(*x*)*}*, the reaction (flip dynamics) is

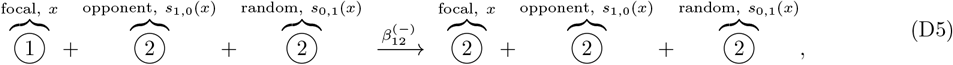

where the flip rate 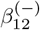 is

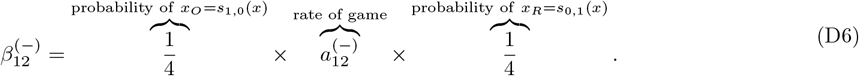

The position of the focal player is *x, i*.*e*., *x*_*F*_ = *x*, the opponent is from (*x*) and it has a different strategy from the focal player, *i.e*., *ξ*(*x*_*O*_) = 2. Suppose the focal player loses a game and gets replaced by an offspring of the opponent, *i.e*., *x*_*R*_ = *x*_*O*_. So, we require *ξ*(*x*) = 1 and *ξ*(*x*_*O*_) = 2. With *x*_*F*_ = *x* fixed, there are 4 feasible combinations of positions {*x*_*F*_, *x*_*O*_, *x*_*R*_}. For every feasible combination, say {*x, s*_1,0_(*x*), *s*_1,0_(*x*)}, the reaction(flip dynamics) is

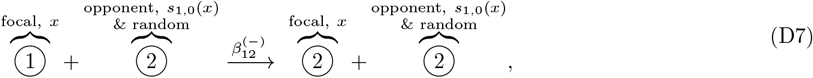

where the flip rate 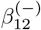 is

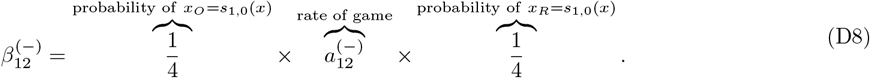

Next, we still assume *ξ*(*x*) = 1, and consider the case where the position of the focal player is *s*_*k, 𝓁*_ (*x*), where |*k*| + |*𝓁* | = 1, *i.e*., the focal player is one of the nearest neighbors of *x*. (Note that there are 4 possible positions of the focal player.) For example, when (*k, ℓ*) = (0, 1), the following three scenarios may result in a state flip on site *x*.

The position of the focal player is *s*_0,1_(*x*), *i.e*., *x*_*F*_ = *s*_0,1_(*x*), and the strategy of the focal player is 2, *i.e*., *ξ*(*x*_*F*_) = 2. The opponent is one of its nearest neighbors (except *x*) and it has the same strategy as *x*_*F*_, *i.e*., *ξ*(*x*_*O*_) = 2. Suppose the focal player wins a game and reproduces and replaces the player at *x* (the third player), *i.e*., *x*_*R*_ = *x*. So, we require *ξ*(*x*_*F*_) = 2, *ξ*(*x*_*O*_) = 2 and *ξ*(*x*) = 1. With *x*_*R*_ = *x* and *x*_*F*_ = *s*_0,1_(*x*) fixed, there are 3 feasible combinations of positions {*x*_*F*_, *x*_*O*_, *x*_*R*_}. For every feasible combination, say {*s*_0,1_(*x*), *s*_1,1_(*x*), *x*}, the reaction (flip dynamics) is

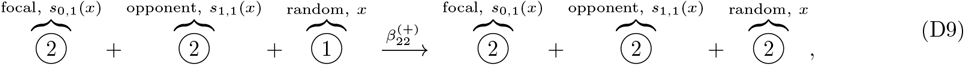

where the flip rate 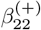 is

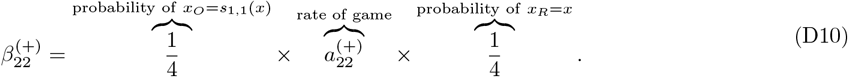

- The position of the focal player is *s*_0,1_(*x*), *i.e*., *x*_*F*_ = *s*_0,1_(*x*), and the strategy of the focal player is 2, *i.e*., *ξ*(*x*_*F*_) = 2. The opponent is one of its nearest neighbor sites (except *x*) and it has a different strategy than *x*_*F*_, *i.e*., *ξ*(*x*_*O*_) = 1. Suppose the focal player wins a game and reproduces and replaces the player at *x* (the third player), *i.e*., *x*_*R*_ = *x*. So, we require *ξ*(*x*_*F*_) = 2, *ξ*(*x*_*O*_) = 1 and *ξ*(*x*) = 1. With *x*_*R*_ = *x* and *x*_*F*_ = *s*_0,1_(*x*) fixed, there are 3 feasible combinations of positions *{x*_*F*_, *x*_*O*_, *x*_*R*_*}*. For every feasible combination, say *{s*_0,1_(*x*), *s*_1,1_(*x*), *x}*, the reaction (flip dynamics) is

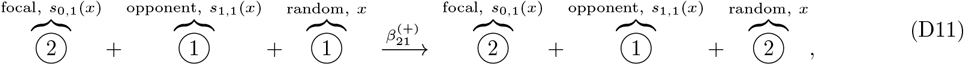

where the flip rate 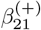 is

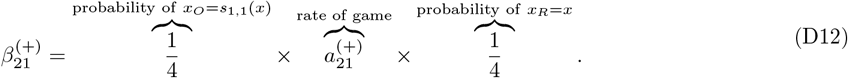
- The position of the focal player is *s*_0,1_(*x*), *i.e*., *x*_*F*_ = *s*_0,1_(*x*), and the strategy of the focal player is 2, *i.e*., *ξ*(*x*_*F*_) = 2. The position of the opponent is *x, i*.*e*., *x*_*O*_ = *x*. Suppose the focal player wins a game and reproduces and replaces the player at *x* (the third player), *i.e*., *x*_*R*_ = *x*. So, we require *ξ*(*x*_*F*_) = 2, *ξ*(*x*) = 1.

With *x*_*O*_ = *x*_*R*_ = *x* and *x*_*F*_ = *s*_0,1_(*x*) fixed, there is only 1 feasible combination of positions {*s*_1,0_(*x*), *x, x*}, and the reaction (flip dynamics) is

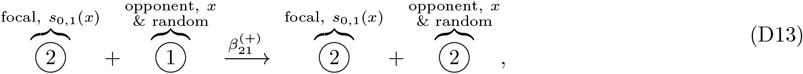

where the flip rate 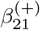 is

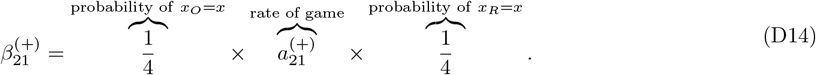

Now, we assume *ξ*(*x*) = 2 and the position of the focal player is *x*, and discuss the following scenarios that may result in a state flip on site *x*. Note that the following scenarios are parallel with the ones that we discussed above with *ξ*(*x*) = 1, except that states 1 and 2 are interchanged.

- The position of the focal player is *x, i*.*e*., *x*_*F*_ = *x*, the opponent is from (*x*) and it has the same strategy as the focal player, *i.e*., *ξ*(*x*_*O*_) = *ξ*(*x*_*F*_) = 2. Suppose the focal player loses a game and gets replaced by a randomly chosen third player in (*x*) (other than the opponent) with a different strategy, *i.e*., *ξ*(*x*_*R*_) = 1. So, we require *ξ*(*x*) = 2, *ξ*(*x*_*O*_) = 2 and *ξ*(*x*_*R*_) = 1. With *x*_*F*_ = *x* fixed, there are 12 feasible combinations of positions *{x*_*F*_, *x*_*O*_, *x*_*R*_*}*. For every feasible combination, say *{x, s*_1,0_(*x*), *s*_0,1_(*x*)*}*, the reaction (flip dynamics) is

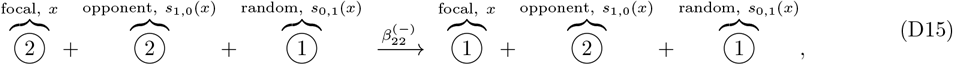

where the flip rate 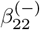 is

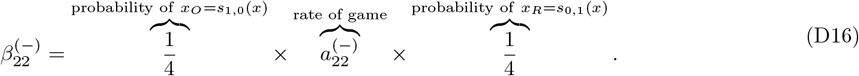
- The position of the focal player is *x, i*.*e*., *x*_*F*_ = *x*, the opponent is from 𝒩 (*x*) and it has a different strategy from the focal player, *i.e*., *ξ*(*x*_*O*_) = 1. Suppose the focal player loses a game and gets replaced by a randomly chosen third player (other than the opponent) with same strategy as the opponent, *i.e*., *ξ*(*x*_*R*_) = 1. So, we require *ξ*(*x*) = 2, *ξ*(*x*_*O*_) = 1 and *ξ*(*x*_*R*_) = 1. With *x*_*F*_ = *x* fixed, there are 12 feasible combinations of positions *{x*_*F*_, *x*_*O*_, *x*_*R*_*}*. For every feasible combination, say *{x, s*_1,0_(*x*), *s*_0,1_(*x*)*}*, the reaction (flip dynamics) is

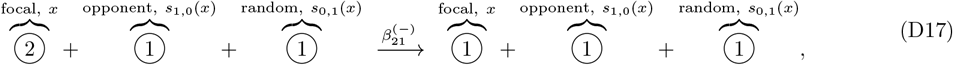

where the flip rate 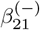 is

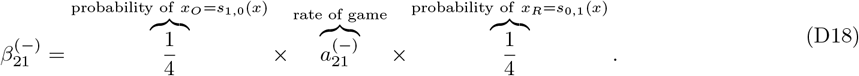

The position of the focal player is *x, i*.*e*., *x*_*F*_ = *x*, the opponent is from (*x*) and it has a different strategy from the focal player, *i.e*., *ξ*(*x*_*O*_) = 1. Suppose the focal player loses a game and gets replaced by an offspring of the opponent, *i.e*., *x*_*R*_ = *x*_*O*_. So, we require *ξ*(*x*) = 2 and *ξ*(*x*_*O*_) = 1. With *x*_*F*_ = *x* fixed, there are 4 feasible combinations of positions {*x*_*F*_, *x*_*O*_, *x*_*R*_}. For every feasible combination, say {*x, s*_1,0_(*x*), *s*_1,0_(*x*)}, the reaction (flip dynamics) is

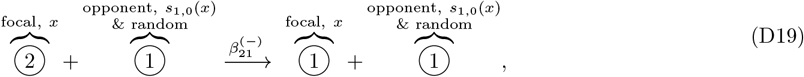

where the flip rate 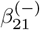 is

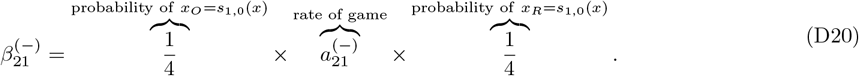

Next, we still assume *ξ*(*x*) = 2, and consider the case where the position of the focal player is *s*_*k,f*_(*x*), where |*k*| + |*𝓁* | = 1, *i.e*., the focal player is one of the nearest neighbors of *x*. (Note that there are 4 possible positions of the focal player.) For example, when (*k, 𝓁*) = (0, 1), the following three scenarios may result in a state flip on site *x*.

The position of the focal player is *s*_0,1_(*x*), *i.e*., *x*_*F*_ = *s*_0,1_(*x*), and the strategy of the focal player is 1, *i.e*., *ξ*(*x*_*F*_) = 1. The opponent is one of its nearest neighbors (except *x*) and it has the same strategy as *x*_*F*_, *i.e*., *ξ*(*x*_*O*_) = 1. Suppose the focal player wins a game and reproduces and replaces the player at *x* (the third player), *i.e*., *x*_*R*_ = *x*. So, we require *ξ*(*x*_*F*_) = 1, *ξ*(*x*_*O*_) = 1 and *ξ*(*x*) = 2. With *x*_*R*_ = *x* and *x*_*F*_ = *s*_0,1_(*x*) fixed, there are 3 feasible combinations of positions {*x*_*F*_, *x*_*O*_, *x*_*R*_}. For every feasible combination, say {*s*_0,1_(*x*), *s*_1,1_(*x*), *x*}, the reaction (flip dynamics) is

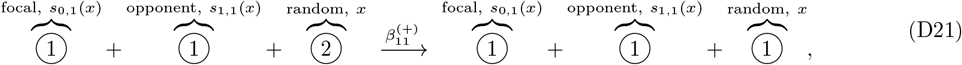

where the flip rate 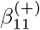 is

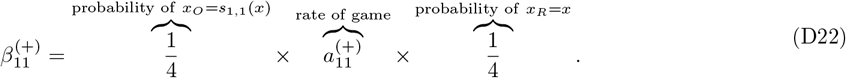

- The position of the focal player is *s*_0,1_(*x*), *i.e*., *x*_*F*_ = *s*_0,1_(*x*), and the strategy of the focal player is 1, *i.e*., *ξ*(*x*_*F*_) = 1. The opponent is one of its nearest neighbor sites (except *x*) and it has a different strategy than *x*_*F*_, *i.e*., *ξ*(*x*_*O*_) = 2. Suppose the focal player wins a game and reproduces and replaces the player at *x* (the third player), *i.e*., *x*_*R*_ = *x*. So, we require *ξ*(*x*_*F*_) = 1, *ξ*(*x*_*O*_) = 2 and *ξ*(*x*) = 2. With *x*_*R*_ = *x* and *x*_*F*_ = *s*_0,1_(*x*) fixed, there are 3 feasible combinations of positions *{x*_*F*_, *x*_*O*_, *x*_*R*_*}*. For every feasible combination, say *{s*_0,1_(*x*), *s*_1,1_(*x*), *x}*, the reaction (flip dynamics) is

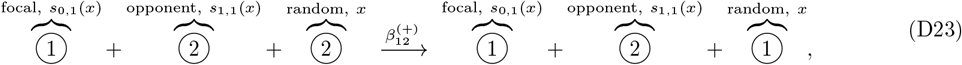

where the flip rate 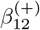 is

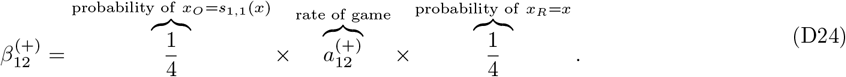

The position of the focal player is *s*_0,1_(*x*), *i.e*., *x*_*F*_ = *s*_0,1_(*x*), and the strategy of the focal player is 1, *i.e*., *ξ*(*x*_*F*_) = 1. The position of the opponent is *x, i*.*e*., *x*_*O*_ = *x*. Suppose the focal player wins a game and reproduces and replaces player at *x* (the third player), *i.e*., *x*_*R*_ = *x*. So, we require *ξ*(*x*_*F*_) = 1, *ξ*(*x*) = 2. With *x*_*O*_ = *x*_*R*_ = *x* and *x*_*F*_ = *s*_0,1_(*x*) fixed, there is only 1 feasible combination of positions {*s*_1,0_(*x*), *x, x*}, and the reaction (flip dynamics) is

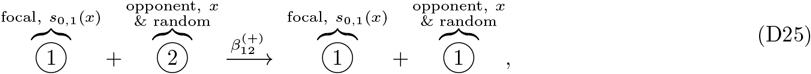

where the flip rate 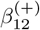 is

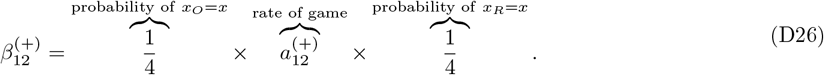

## Appendix E

**Hydrodynamic limit of spatial, stochastic games**

In addition to the partial flip dynamics described above, we introduce another type of dynamics, namely the particle exchange dynamics. Consider the scaled lattice 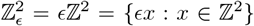. Assume the states at sites *x* and *y* are exchanged at rate *κ ϵ*^*−*2^, where ‖ *x* − *y* ‖ _1_ = *ϵ* and *κ* is the *diffusion constant*. This is equivalent to say that the player at position *x* tries to swap with another nearest player (chosen at random with equal transition probability 1*/*4) at rate 2*κ ϵ*^*−*2^. Specifically, if the rate of *x, y* swapping (given *x* is the center and *y* is the neighbor) is 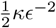, the same swap event could also occur in the case of *y* as the center and *x* as the neighbor with swap rate 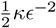. Therefore, the total rate of *x, y* swapping is *κ* ϵ^*−*2^. The configuration of the scaled system at time *t* is denoted by 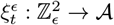.

In an appropriately scaled limit, in which the lattice spacing goes to zero (ϵ→ 0) and the speed of stirring goes to infinity (ϵ^*−*2^→ ∞), the densities of the different types of particles converge to continuous densities which are solutions of a reaction-diffusion equation [23, 37, 38]. The term ‘reaction’ corresponds to the particle flip dynamics and the term ‘diffusion’ corresponds to the scaled particle exchange dynamics (fast stirring). In the next theorem, we will derive the reaction-diffusion equation as the hydrodynamic limit of the interacting particle system, where the particle exchange dynamics is as above, and the particle flip dynamics is as described in Appendix D.

### Theorem 1.

*Consider an interacting particle system on the scaled lattice* ℤ^2^. *Let the particle flip dynamics be given in Eqs. D3 - D26, and the particle exchange dynamics are given by the symmetric nearest neighbor stirring with scaled rate* 2*κ ϵ*^*−*2^. *Suppose* 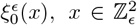, *are independent (product measure) and let* 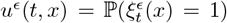. *If u*^*ϵ*^ (0, *x*) = *g*(*x*) *is continuous then as ϵ* → 0, *u*^*ϵ*^ (*t, x*) *converges to the hydrodynamic limit u*(*t, x*), *where u*(*t, x*) *is the bounded solution of*

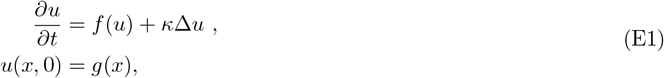

*where the reaction term f* (*u*) *is*

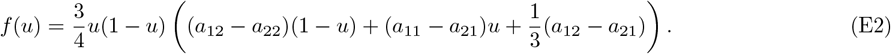

### Proof

According to theory of hydrodynamic limits of interacting particle systems [23, 37, 38], the reaction term *f* (*u*) is

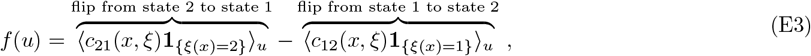

where ⟨. ⟩ _*u*_ denotes the expected value under product measure in which states 1 and 2 have densities *u* and 1 *− u* respectively, *i.e*., ℙ (*ξ*(*x*) = 1) = *u* and ℙ (*ξ*(*x*) = 2) = 1 *− u*. Recall the flip dynamics that are associated with *ξ*(*x*) = 2, Eqs. D15 - D26, the first term *c*_21_(*x, ξ*)**1**_*{ξ*(*x*)=2*} u*_ becomes

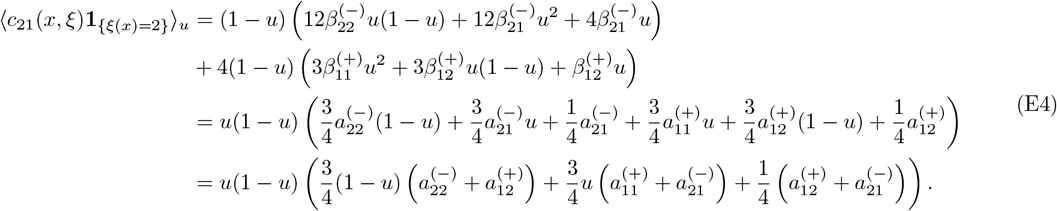

Similarly, we let *ξ*(*x*) = 1, using Eqs. D3 - D14, the second term *c*_12_(*x, ξ*)**1**_*{ξ*(*x*)=1*} u*_ becomes

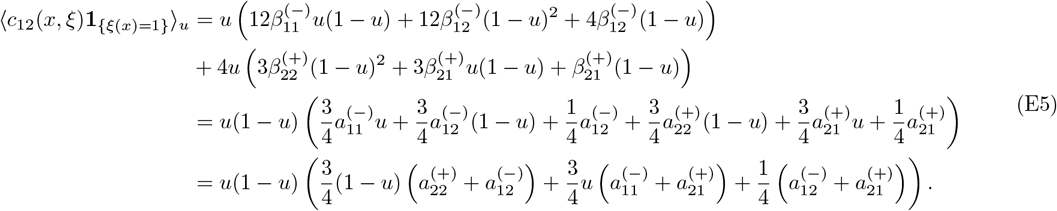

Plugging Eqs. E4 - E5 into Eq. E3, we have

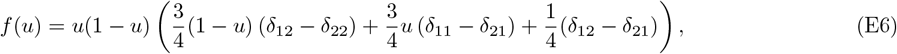

where 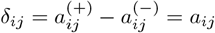.

### Remark 1.

*Comparing Eqs. E1–E2 with Eq. 2, we point out that the limiting reaction-diffusion equation associated with the spatial game system is different from simply adding a diffusion term to the mean-field replicator dynamics. This observation recapitulates the findings in [24]. This is because for each spatial game in dimension 2, there is a 1/4 probability that the random third player coincides with the opponent, which is not negligible. In contrast, in a non-spatial (mean-field) system the probability would be* 1*/N (here N is the total number of players), which goes to zero as N* → ∞. *In addition, in dimension d, a similar derivation leads to a reaction term*

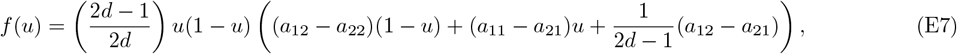

*which recovers the replicator dynamics when d is large*, 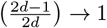 *as d* ≫ 1.

*On the other hand, we can modify the game rules such that the opponent player cannot be identified as the random selected third player in Eqs. D3 - D26, i*.*e*., *we guarantee the random selected third player is not an opponent, then the hydrodynamics PDE takes the form of Eq. E1 with reaction term Eq. A11*.

The reaction term *f* (*u*) in Eq. E2 is a cubic polynomial. Let us define the potential function *W* associated with *f* as

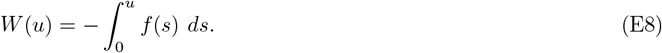

If *W* has two local minima at *u* = 0 and *u* = 1, and a local maximum at *u* = *θ* where *θ* ∈ (0, 1), then we refer to *W* as a *double-well potential*. In the following proposition, we give a sufficient condition for *W* to be a double-well potential.

### Proposition E.1.

*For the spatial game in dimension d, suppose the* 2 *×* 2 *payoff matrix A satisfies*

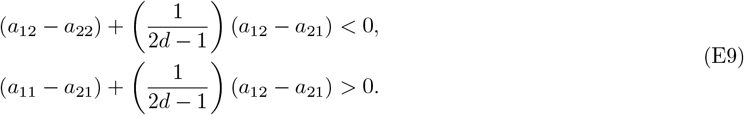

*Then, the potential function W* (*u*) *in Eq. E8 is a double-well potential*.

### Proof

To ensure the potential function *W* has a double-well shape, we restrict *f* (*u*) = 0 when *u* = 0, 1 and *θ*, where *θ* ∈ (0, 1), and *f* (0) *<* 0, *f* (1) *<* 0, *f* (*θ*) *>* 0. Note that *f* (*u*) = 0 whenever *u* = 0, 1 given the form of *f* (*u*) in Eq. E7. Define a linear function *h* on [0, 1] as

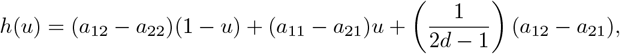

so that 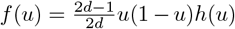. To have *f* (*θ*) = 0 with *θ* ∈ (0, 1), we require *h*(*θ*) = 0 for some *θ* ∈ (0, 1). Since *h* is linear, we need sign*{h*(0)*}* ≠ sign*{h*(1)*}* to ensure the existence of such a *θ*, moreover, we need *h*(0) *<* 0 and *h*(1) *>* 0 so that *f* (*θ*) *>* 0. In doing so, we obtain Eq. E9.

## Decay of free energy

The reaction-diffusion equation in Theorem 1 takes the form of an Allen-Cahn equation [16]. When Eq. E1 is defined on a two dimensional torus 𝕋^2^, the evolution of Eq. E1 can be viewed as the *L*^2^-gradient flow of the Ginzburg-Landau free energy functional [25, 26]

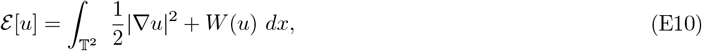

*i.e*., 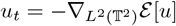. Taking the derivative of ℰ [*u*] with respect to time yields

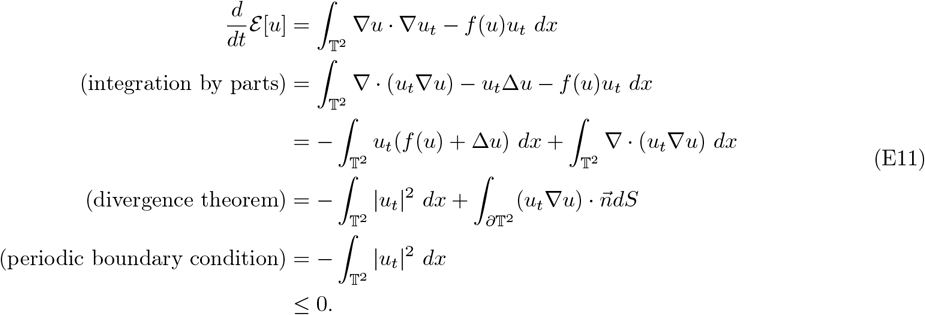

Therefore, the free energy is non-increasing in time. Note that the free energy ℰ [*u*] contains two terms, where the gradient term penalizes spatial variation (thus has a smoothing effect), whereas the potential *W* (which has a double-well shape with two local minima) drives the system to undergo phase separation.

## Linear stability analysis

For Eq. E1, note that the constant solution *u*(*x, t*) ≡ *ū* is a steady state, where *ū* ∈ (0, 1) is such that *f* (*ū*) = 0, and it corresponds to a mixed strategy that is spatially homogeneous. Now let us consider a solution that has a small fluctuation around *ū*, that is, *u*(*x, t*) = *ū* + *du*(*x, t*), where *du* |≪ | 1. Substituting this solution to the evolution equation Eq. E1 and omitting the terms of order (*δu*)^2^ or higher, we find that an arbitrarily small perturbation *δu* evolves as follows:

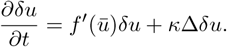

The general solution of a constant-coefficient linear evolution equation can be written as *δu*(*x, t*) = *ve*^*σt*^*e*^*iqx*^, where *v* is some nonzero constant vector, *σ* is growth rate and *q* = [*kπ/ℓ, jπ/ℓ*]^*T*^ is the wave vector (*qx* is a dot product). Substituting *δu*(*x, t*) = *ve*^*σt*^*e*^*iqx*^ into the above linear evolution equation and canceling common terms from both sides, we find that

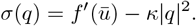

The growth rate *σ* is then determined as a function of the wave vector. Those associated wavelengths with positive growth rates are unstable, and those with negative growth rates are stable; the parameters of the system determine which, if any, of the wavelengths fall into each category. Note that we have *f* (*ū*) *>* 0 when *W* is a double-well potential, which leads to *σ*(0) *>* 0. In addition, if either *κ* is sufficiently small or the domain is sufficiently large, we also have *σ*(*q*) *>* 0 for all |*q*| small, indicating that a small fluctuation could lead to phase diverge from the homogeneous equilibrium.

